# Syntaxin-1A modulates vesicle fusion in mammalian neurons via juxtamembrane domain dependent palmitoylation of its transmembrane domain

**DOI:** 10.1101/2022.03.11.483937

**Authors:** Gülçin Vardar, Andrea Salazar-Lázaro, Sina Zobel, Thorsten Trimbuch, Christian Rosenmund

**Affiliations:** Department of Neurophysiology, Charité-Universitätsmedizin Berlin, Humboldt-Universität zu Berlin, and Berlin Institute of Health, 10117 Berlin, Germany; NeuroCure Excellence Cluster, Berlin, Germany

**Keywords:** juxtamembrane domain, transmembrane domain, Syntaxin-1 (STX1), palmitoylation

## Abstract

SNAREs are undoubtedly one of the core elements of synaptic transmission. On the contrary to the well characterized function of their SNARE domains bringing the plasma and vesicular membranes together, the level of contribution of their juxtamembrane domain (JMD) and the transmembrane domain (TMD) to the vesicle fusion is still under debate. To elucidate this issue, we analyzed three groups of STX1A mutations: 1) elongation of STX1A’s JMD by three amino acid insertions in the junction of SNARE-JMD or JMD-TMD; 2) charge reversal mutations in STX1A’s JMD; and 3) palmitoylation deficiency mutations in STX1A’s TMD. We found that both JMD elongations and charge reversal mutations have position dependent differential effects on Ca^2+^-evoked and spontaneous neurotransmitter release in cultured murine hippocampal neurons. Importantly, we show that STX1A’s JMD regulates the palmitoylation of STX1A’s TMD and loss of STX1A palmitoylation either through charge reversal mutation K260E or by loss of TMD cysteines particularly inhibits spontaneous vesicle fusion. Interestingly, the retinal ribbon specific STX3B has a glutamate in the position corresponding to the K260E mutation in STX1A and mutating it with E259K acts as a molecular on-switch. Furthermore, palmitoylation of post-synaptic STX3A can be induced by the exchange of its JMD with STX1A’s JMD together with the incorporation of two cysteines into its TMD. Forced palmitoylation of STX3A dramatically enhances spontaneous vesicle fusion suggesting that STX1A regulates spontaneous release through two distinct mechanisms: one through the C-terminal half of its SNARE domain and the other through the palmitoylation of its TMD.

## Introduction

Numerous intracellular trafficking pathways utilize vesicle fusion of various types for that the SNARE proteins play a pivotal role. Similarly, synaptic transmission as a means of neuronal communication employs the fusion of the neurotransmitter containing synaptic vesicles (SVs) with the presynaptic plasma membrane. Therefore, presynaptic vesicular release, too, largely relies on the SNARE complex in this case formed by the presynaptic neuronal SNAREs synaptobrevin-2 (SYB2), SNAP-25 and syntaxin-1A or syntaxin-1B (STX1), assisted by modulatory synaptic proteins (Rizo, 2018).

Both STX1 and SYB2 are integral proteins on the plasma and vesicular membranes, respectively, and their C-terminal transmembrane domain (TMD) and SNARE motif are separated only by a short polybasic juxtamembrane domain (JMD). By contrast, SNAP-25 is anchored to the plasma membrane through the palmitoylated cysteines in its linker region between its two SNARE motifs. The N-to-C helical formation of the *trans*-SNARE complex leads to the apposition of the two membranes (Gao et al., 2012; Sorensen et al., 2006; Stein et al., 2009) in preparation for membrane merger. Additionally, the formation of the *cis*-SNARE complex on the plasma membrane after vesicle fusion suggests that the SNAREs further zipper into the JMD and TMDs of STX1 and SYB2 (Hernandez et al., 2012; Risselada et al., 2011; Stein et al., 2009).

Importantly, the merger of two lipid membranes *per se* is an energetically high-cost process (Risselada and Mayer, 2020; Zhang, 2017) where the reduction of the energy barrier for membrane merger has been primarily assigned to the modulatory proteins, such as synaptotagmin-1 (SYT1) (Hui et al., 2009; Martens et al., 2007). Whereas the role of the SNARE domain zippering is well characterized in vesicle fusion, the roles of TMD and JMDs of STX1 and SYB2 is poorly understood as it is still under dispute whether they are active or rather superfluous factors in that process (Han et al., 2017).

At this point, it is critical to note that the assumption of the TMDs of STX1 and SYB2 as a passive membrane anchor (Zhou et al., 2013) is not compatible with their presumed β-branched nature, which confers a high TMD flexibility (Dhara et al., 2016; Hastoy et al., 2017) and which is uncommon among integral transmembrane proteins (Quint et al., 2010). Remarkably, another feature shared by STX1 and SYB2 but unusual for the majority of transmembrane proteins is that they are palmitoylated in their TMDs (Kang et al., 2008; Prescott et al., 2009) which is shown as an alternative mechanism for TMD tilting and flexibility in the membrane (Blaskovic et al., 2013; Charollais and Van Der Goot, 2009). Whichever mechanism underlies the oblique position of the TMDs of STX1 and SYB2 in the membrane, it is known that their tilted nature causes their polybasic JMDs to be immersed into the membrane potentially neutralizing the repulsive forces between the apposed vesicular and plasma membrane (Kim et al., 2002; Kweon et al., 2002; Williams et al., 2009). Conceivably therefore, the JMD and TMD of STX1 and SYB2 might actively regulate vesicle fusion by reducing the energy barrier for membrane merger.

Based on this dispute, we addressed whether STX1A play additional roles in vesicle fusion to facilitate the membrane merger through its JMD and TMD and created STX1A mutants in which its JMD is elongated or modified by charge reversal single point mutations at different positions that lead to altered electrostatic nature. We also constructed palmitoylation deficient STX1A mutants and analyzed electrophysiological properties of all STX1A mutants in STX1-null hippocampal mouse neurons. Firstly, we found that the tight coupling of STX1A’s JMD to its SNARE domain but not to its TMD is fundamental for neurotransmitter release. Secondly and strikingly, we found that the palmitoylation of STX1A’s TMD depends on its JMD’s cationic amino acids and that loss of palmitoylation dramatically impairs spontaneous release while leaving Ca^2+^-evoked release almost intact. Furthermore, we successfully emulated the regulation of palmitoylation of STX1A’s TMD by its JMD and its effect on neurotransmitter release by using other syntaxin isoforms, STX3A or retinal ribbon specific STX3B (Curtis et al., 2010; Curtis et al., 2008). Based on our data, we propose a direct involvement of STX1A’s JMD and TMD in vesicle fusion through electrostatic forces and palmitoylation.

## Results

### The level and mode of impairment of neurotransmitter release by the elongation of STX1A’s JMD depends on the position of the GSG-insertion

The progressive N-to-C zippering of the *trans*-SNARE complex formed by STX1, SNAP25, and SYB2 sets up the vesicular and plasma membrane in close proximity for vesicle fusion (Gao et al., 2012; Sorensen et al., 2006; Stein et al., 2009). To test how the continuity of the SNARE TMD assembly affects the vesicle fusion, we elongated the JMD of STX1A by an insertion of three amino acids, GSG, leading to an extra helical turn either at the junction of its SNARE domain and JMD (STX1A^GSG259^) or of its JMD and TMD (STX1A^GSG265^) (Figure 1A). Using our lentiviral expression system in STX1-null neurons (Vardar et al., 2016; Vardar et al., 2021) and electrophysiological assessment, we surprisingly found that the insertion of one extra helical turn into the JMD of STX1A led to position specific physiological phenotypes (Figure 1).

**Figure 1:**
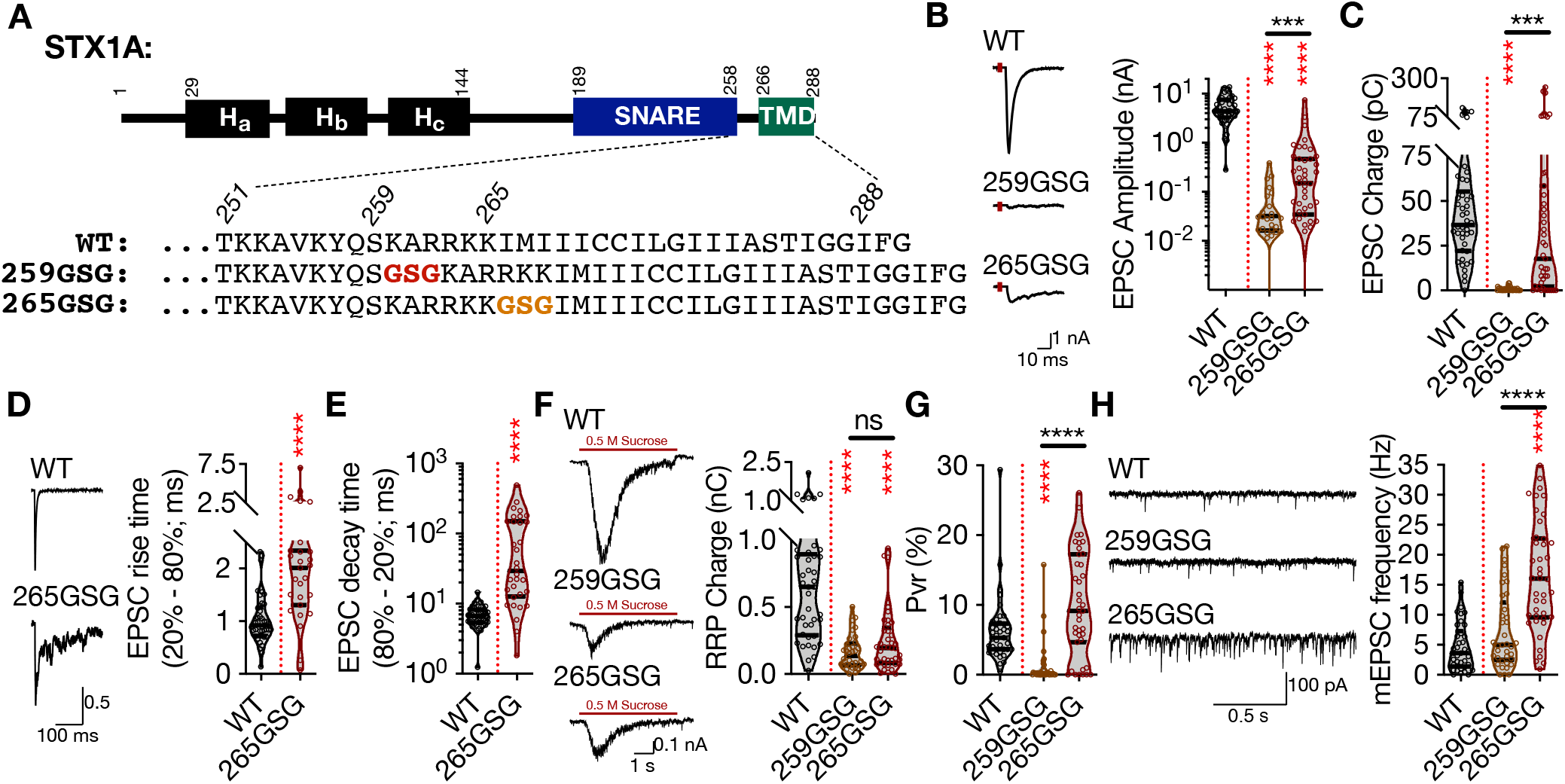
The level and mode of impairment of neurotransmitter release by the elongation of STX1A’s JMD depends the position of the GSG-insertion. A. STX1A domain structure and insertion of GSG into its JMD. The C-terminal TMD and SNARE motif are separated only by a short polybasic JMD. B. Example traces (left) and quantification of the amplitude (right) of EPSCs obtained from hippocampal autaptic STX1-null neurons rescued either with STX1A^WT^, STX1A^GSG259^, or STX1A^GSG265^. C. Quantification of the charge (right) of EPSCs obtained from the same neurons as in (B). D. Example traces with the peak normalized to 1 (left) and quantification of the EPSC rise time measured from 20% to the 80% of the EPSC recorded from STX1B^WT^ or STX1A^GSG265^. E. Quantification of the decay time measured from 80% to the 20% of the EPSC recorded from the same neurons as in (D). F. Example traces (left) and quantification of the charge transfer (right) of 500 mM sucrose-elicited RRPs obtained from the same neurons as in B. G. Quantification of Pvr determined as the percentage of the RRP released upon one AP. H. Example traces (left) and quantification of the frequency (right) of mEPSCs recorded at −70 mV. Data information: The artifacts are blanked in example traces in (B, D, and F). The example traces in (H) were filtered at 1 kHz. In (B–H), data points represent single observations, the violin bars represent the distribution of the data with lines showing the median and the quartiles. Red and black annotations (stars and ns) on the graphs show the significance comparisons to STX1A^WT^ and STX1A^GSG259^, respectively. Nonparametric Kruskal-Wallis test followed by Dunn’s *post hoc* test was applied to data in (B, C, and F-H), Mann-Whitney test was applied in (D & E); ***p ≤ 0.001, ****p ≤ 0.0001. The numerical values are summarized in source data.

Firstly, the amplitude of the excitatory postsynaptic current (EPSC) was reduced to almost zero in both mutants (Figure 1B) suggesting that the force transfer from the SNARE complex formation to the merging of the plasma and vesicular membranes is strictly regulated by the length of the JMD of STX1A. On the contrary to the EPSC amplitude, the EPSC charge analysis revealed that STX1A^GSG259^ and STX1A^GSG265^ had differential effects on the vesicle fusion *per se* (Figure 1C). Whereas STX1A^GSG259^ completely blocked Ca^2+^-evoked release, STX1A^GSG265^ only slowed it down as expressed by a 2-fold and more than 10-fold increase in the EPSC rise and the decay time, respectively (Figure 1D and 1E). This suggests that decoupling STX1A’s JMD from its SNARE domain has more deleterious effects on Ca^2+^-evoked neurotransmitter release than decoupling it from its TMD has. Interestingly, elongation of STX1A’s JMD at either position impaired not only Ca^2+^-evoked neurotransmitter release but also the upstream process, vesicle priming, as shown by a significant decrease in the size of the readily releasable pool (RRP) of SVs to ~ 25% by STX1A^GSG259^ and to ~ 40% by STX1A^GSG265^ (Figure 1F), consistent with previous studies (Zhou et al., 2013). This led to an abolishment of vesicular probability (Pvr) in the STX1A^GSG259^ neurons and a trend towards an increase in Pvr in the STX1A^GSG265^ neurons (Figure 1G).

Once again, three amino acid insertion between the SNARE domain and the TMD of STX1A had different effects on spontaneous neurotransmitter release depending on the position of the GSG-insertion. Whereas uncoupling of the SNARE domain and the TMD of STX1A by GSG259 mutation had no effect on spontaneous neurotransmission, uncoupling of its JMD and the TMD by GSG265 mutation increased the miniature EPSC (mEPSC) frequency by ~3-fold compared to that of STX1A^WT^ (Figure 1H). This is interesting as it shows that it is not only the length of the linker region but also the interplay between the SNARE domain-JMD and JMD-TMD that is important for the regulation of synchronous Ca^2+^-evoked and spontaneous release.

### Charge reversal mutations in STX1A’s JMD manifest position specific effects on different modes of neurotransmitter release and on molecular weight of STX1A

JMD of STX1A, ‘260-KARRKK-265’, consists of basic residues with characteristics of PIP2 binding motif (van den Bogaart et al., 2011a). In fact, it has been shown that it drives PIP2 or PIP3 dependent clustering of STX1A (Khuong et al., 2013; van den Bogaart et al., 2011a) and the inhibition of its interaction with PIP2/PIP3 leads to defects in neurotransmitter release (Khuong et al., 2013). Additionally, the JMD of STX1A is embedded in the membrane due to the tilted conformation of STX1A’s TMD (Kim et al., 2002; Kweon et al., 2002) setting a ground for a possible role in vesicle fusion through electrostatic interactions with the plasma membrane. Therefore, it is plausible that the position dependent differential effects of the GSG-insertion might be a result of differential perturbations in JMD-membrane interactions. To test this, we introduced single amino acid charge reversal mutations into the STX1A’s JMD (Figure 2 and Figure 2-Supplement 1). For that purpose, we mutated the lysine or arginine residues from amino acid (aa) 256 to aa 265 into glutamate to achieve maximum alterations in the electrostatic interactions between the JMD and the plasma membrane, where K256E served as a control for the effects of net charge difference only (Figure 2A).

**Figure 2:**
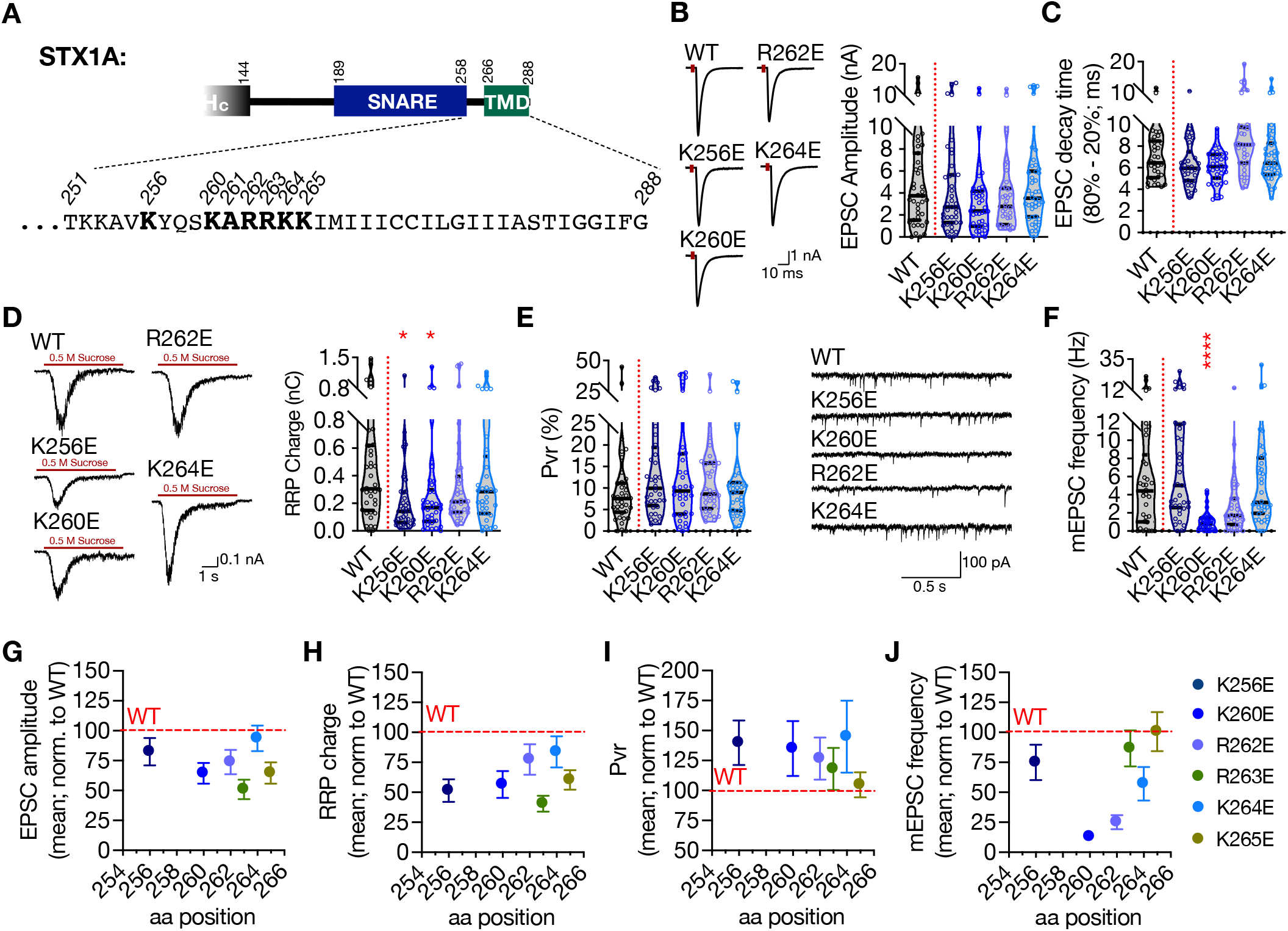
Charge reversal mutations in STX1A’s JMD manifest position specific effects on different modes of neurotransmitter release. A. Position of the charge reversal mutations on STX1A’s JMD. B. Example traces (left) and quantification of the amplitude (right) of EPSCs obtained from hippocampal autaptic STX1-null neurons rescued either with STX1A^WT^, STX1A^K256E^, STX1A^K260E^, STX1A^R262E^, or STX1A^K264E^. C. Quantification of the decay time (80% - 20%) of the EPSC recorded from the same neurons as in (B). D. Example traces (left) and quantification of RRP recorded from the same neurons as in (B). E. Quantification of Pvr recorded from the same neurons as in (B). F. Example traces (left) and quantification of the frequency (right) of mEPSCs recorded from the same neurons as in (B). G. Correlation of the EPSC amplitude normalized to that of STX1A^WT^ to the position of the charge reversal mutation. H. Correlation of the RRP charge normalized to that of STX1A^WT^ to the position of the charge reversal mutation. I. Correlation of Pvr normalized to that of STX1A^WT^ to the position of the charge reversal mutation. J. Correlation of the mEPSC frequency normalized to that of STX1A^WT^ to the position of the charge reversal mutation. Data information: The artifacts are blanked in example traces in (B) and (D). The example traces in (F) were filtered at 1 kHz. In (B–F), data points represent single observations, the violin bars represent the distribution of the data with lines showing the median and the quartiles. In (G-J), data points represent mean ± SEM. Red annotations (stars) on the graphs show the significance comparisons to STX1A^WT^. Nonparametric Kruskal-Wallis test followed by Dunn’s *post hoc* test was applied to data in (B-F); *p ≤ 0.05, ***p ≤ 0.001, ****p ≤ 0.0001. The numerical values are summarized in source data.

Unlike the GSG insertion mutants, charge reversal mutations did not lead to major changes in the EPSC size and kinetics (Figure 2B, 2C, and Figure 2 - Supplement 1). Overall, whereas STX1A^R263E^ significantly decreased the EPSC amplitude to ~ 50% compared to that of STX1A^WT^, the other charge reversal mutants only showed a trend towards a 10-35% decrease in EPSC amplitude (Figure 2B and Figure 2 - Supplement 1). Similarly, not all the charge reversal mutants led to an impairment in vesicle priming but only STX1A^K256E^, STX1A^K260E^ and STX1A^R263E^ mutants reduced the RRP size (Figure 2D and Figure 2 - Supplement 1). This suggests that the observed impairments in the Ca^2+^-evoked release and vesicle priming are not simply due to the change of the net total charge of STX1A’s JMD and that the role of its electrostatic interactions with the plasma membrane are position dependent. Additionally, none of the mutants altered the Pvr (Figure 2E and Figure 2 - Supplement 1).

Interestingly, however, charge reversal mutants in STX1A’s JMD led to differential results in the spontaneous release. Most of the mutants had no effect on the mEPSC frequency and STX1A^R262E^ showed only a trend towards a decrease by ~50% with a p-value of 0.46 compared to that of STX1A^WT^ (Figure 2F and Figure 2 - Supplement 1). However, STX1A^K260E^ mutant unexpectedly showed a dramatic and significant decrease in spontaneous release as it reduced the mEPSC frequency by ~80% from 5.6 Hz to 1 Hz (Figure 2F). This again suggests that the perturbation of the electrostatic interactions between the plasma membrane and the JMD of STX1A affect the neurotransmitter release in a position specific manner. For a better visualization of the position specific effects of the charge reversal mutations on release parameters, we plotted the EPSC amplitude, RRP size, Pvr, and mEPSC frequency values normalized to the values obtained from STX1A as a function of the aa position of the charge reversal mutations (Figure 2G – 2J). Whereas the alterations in the EPSC amplitude, RRP size, and Pvr showed no correlation to the position of the basic to acidic mutations (Figure 2G–2I), spontaneous release approved to be specifically perturbed by glutamate insertion at the N-terminus of JMD, that is STX1A^K260E^ (Figure 2J). Importantly, K256E mutation which resides in the SNARE domain and thus more N-terminally to the JMD showed no impact on the spontaneous release suggesting a specific function for STX1A’s JMD in the regulation of spontaneous neurotransmitter release (Figure 2F and 2J).

Both the SNARE domain and the TMD of STX1A have helical structures whereas its JMD has an unstructured nature (Kim et al., 2002; Kweon et al., 2002). This might potentially create a helix-loop-helix formation between the SNARE domain and the TMD when STX1A is isolated. It is known that the mutations potentially affecting the helix-loop-helix formation in the membrane proteins can alter their electrophoresis speed in an SDS-PAGE gel (Rath et al., 2009). To test how the charge reversal mutations on STX1A’s JMD affect the electrophoretic behavior of STX1A, we probed our mutants on Western Blot (WB) using neuronal lysates obtained from high-density cultures (Figure 3A). Surprisingly, STX1A’s JMD charge reversal mutations caused not only an apparent different molecular weight on the SDS-PAGE, but they also showed differing band patterns, with the addition of two lower band sizes (Figure 3A). To analyze the effect of the STX1A’s JMD charge reversal mutations on STX1A’s SDS-PAGE behavior, we assigned arbitrary hierarchical numbers from 1 to 6 based on the distance traveled and the number of bands as visualized by STX1A antibody, where number 1 represents the lowest single band as in STX1A^K260E^ and number 6 represents the highest single band as in STX1A^WT^ (Figure 3B). We also plotted the weighed band intensity for STX1A for which the lowest band was arbitrarily assigned by 1X and the highest band was assigned as 100X as to hierarchically measure the intensity of the bands (Figure 3C). We observed a correlation between the SDS-PAGE behavior of STX1A to the position of the charge reversal mutations on its JMD with K260E mutation causing the most dramatic change in the WB band pattern (Figure 3B and 3C). To test whether the change in the WB band pattern of STX1A lysates is only due to the differential velocity of lysine and glutamate in an SDS-PAGE or whether it reflects the presence or absence of a post-translational modification (PTM), we prepared lysates of HEK293 cell cultures transfected with STX1A^WT^ and charge reversal mutants (Figure 3D). Strikingly, none of the constructs showed a comparable pattern to the STX1A obtained from neurons and the band pattern of all STX1A JMD mutants as well as STX1A^WT^ collapsed to the level of STX1A^K260E^ (Figure 3D – 3F). This suggests that STX1A is post-translationally modified in neurons and this PTM is absent in HEK293 cells. Importantly, it also implies that this PTM is regulated by the basic amino acid residues in the JMD of STX1A in a position dependent manner. To test whether the possible PTM on STX1A correlates with the neurotransmitter release properties, we plotted the EPSC amplitude, RRP size, Pvr, and the mEPSC frequency values normalized to that of STX1A^WT^ as a function of WB band distribution of STX1A^WT^ and charge reversal mutants (Figure 3G – 3J). We observed again that the lower bands specifically represent an impairment in spontaneous neurotransmitter release (Figure 3J).

**Figure 3:**
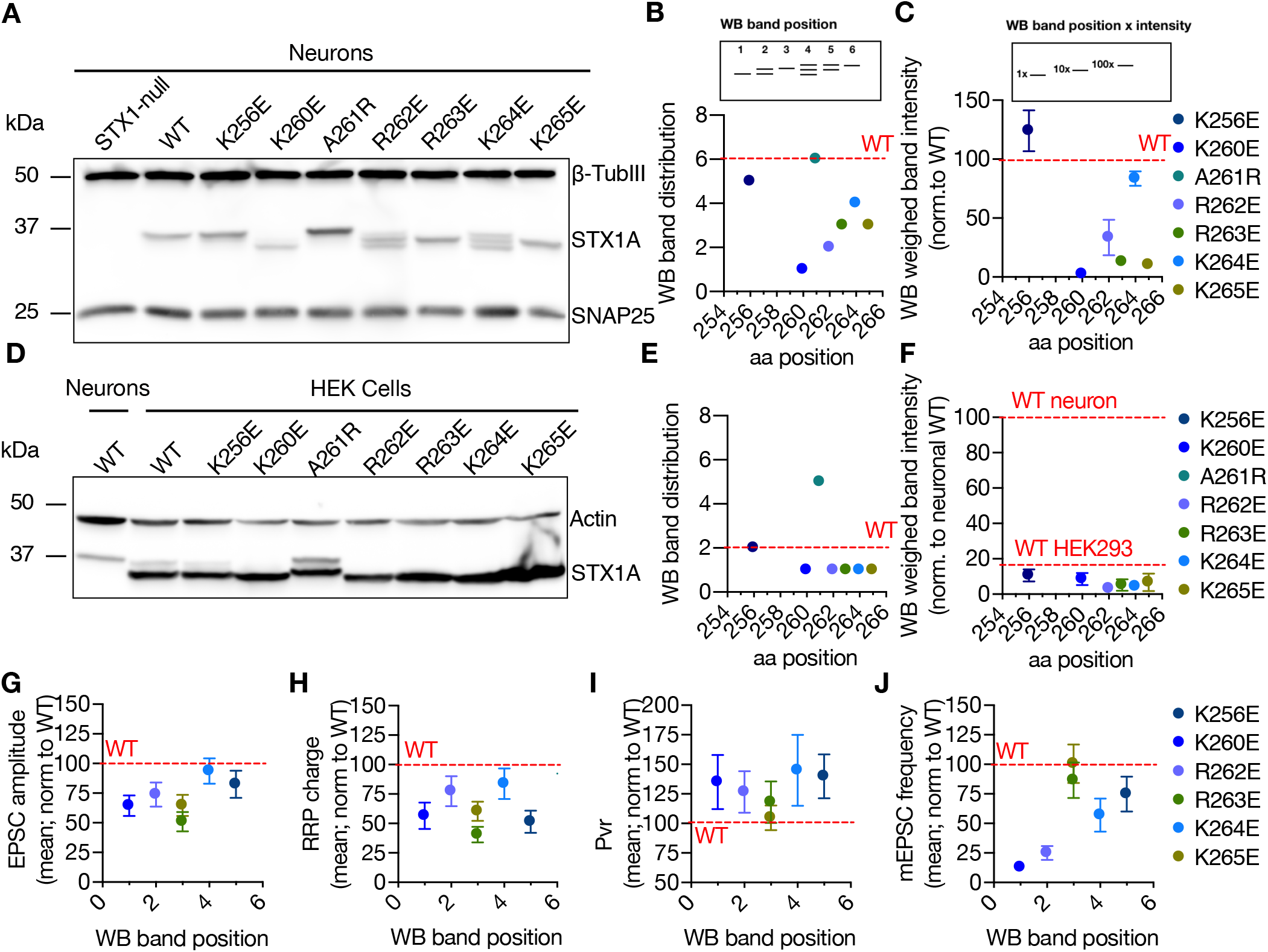
Charge reversal mutations in STX1A’s JMD manifest position specific effects on the molecular weight of STX1A. A. Example image of SDS-PAGE of the electrophoretic analysis of lysates obtained from STX1-null neurons transduced with different STX1A JMD charge reversal mutations. B. Quantification of the STX1A band pattern on SDS-PAGE of neuronal lysates through assignment of arbitrary hierarchical numbers from 1 to 6 based on the distance traveled, where number 1 represents the lowest single band as in STX1A^K260E^ and number 6 represents the highest single band as in STX1A^WT^. C. Quantification of the STX1A weighed band intensity of neuronal lysates for which the lowest band was arbitrarily assigned by 1X and the highest band was assigned as 100X and multiplied by the measured intensity of the STX1A bands. D. Example image of SDS-PAGE of the electrophoretic analysis of lysates obtained from HEK293 cells transfected with different STX1A JMD charge reversal mutations. E. Quantification of the STX1A band pattern on SDS-PAGE of HEK293 cell lysates as in (B). F. Quantification of the STX1A weighed band intensity on SDS-PAGE of HEK293 cell lysates as in (C). G. Correlation of the EPSC amplitude normalized to that of STX1A^WT^ to the WB band position of STX1A charge reversal mutation. H. Correlation of the RRP charge normalized to that of STX1A^WT^ to the WB band position of STX1A charge reversal mutation. I. Correlation of Pvr normalized to that of STX1A^WT^ to the WB band position of STX1A charge reversal mutation. J. Correlation of the mEPSC frequency normalized to that of STX1A^WT^ to the WB band position of STX1A charge reversal mutation. Data information: In (C, F, and G–J), data points represent mean ± SEM. Red lines in all graphs represent the STX1A^WT^ level. The numerical values are summarized in source data.

### STX1A’s JMD modifies the palmitoylation of its TMD which in turn regulates the spontaneous neurotransmitter release

As STX1A charge reversal mutants showed a specific banding pattern on SDS-PAGE when the lysates were obtained from neurons but not from HEK293 cells, we continued with our experiments addressing a potential neuronal specific PTM of STX1A. A modulatory function for a polybasic stretch has been previously shown as a prerequisite for Rac1 palmitoylation (Navarro-Lerida et al., 2012). As STX1A is known to be palmitoylated on its two cysteine residues C271 and C272 in its TMD (Kang et al., 2008) that neighbors its polybasic JMD, we first tested whether the molecular size shift in STX1A^K260E^ is due to loss of palmitoylation. For that purpose, we created either single point mutants STX1A^C271V^ or STX1A^C272V^ or a double point mutant STX1A^CC271,272VV^ (STX1A^CVCV^) and probed them on SDS-PAGE in comparison to STX1A^WT^ and STX1A^K260E^ (Figure 4A and 4B). Whereas STX1A^CVCV^ mutant migrated with an electrophoretic speed corresponding to the size of STX1A^K260E^, the single point mutants STX1A^C271V^ and STX1A^C272V^ showed a molecular size in the middle between STX1A^WT^ and STX1A^K260E^ (Figure 4B). This suggests that the size shift in the charge reversal mutants (Figure 3A) might indeed be due to the impairments in palmitoylation of either cysteine residues (middle bands) or both (lowest band). As a control, we also created a mutant in which CVCV and K260E mutants were combined (STX1A^K260E+CVCV^) and observed that the charge reversal mutation did not cause any further size shift in STX1A^CVCV^ (Figure 4B).

**Figure 4:**
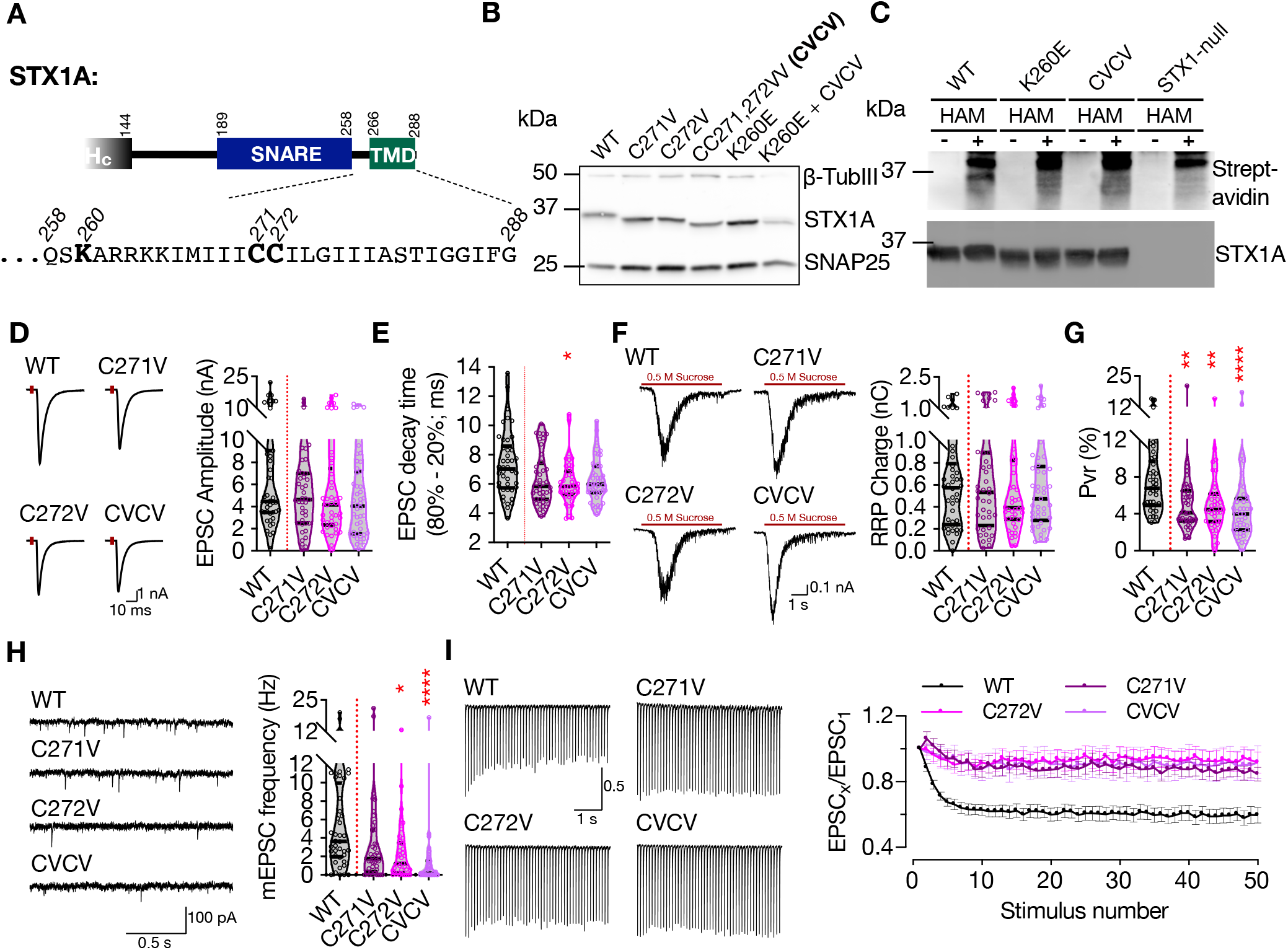
Charge reversal mutations in STX1A’s JMD manifest position specific effects on different modes of neurotransmitter release. A. Position of the palmitoylation deficiency mutations on STX1A’s TMD. B. Example image of SDS-PAGE of the electrophoretic analysis of lysates obtained from STX1-null neurons transduced with STX1A with different palmitoylation deficiency mutations and with STX1AK260E and ^STX1AK260E+CVCV^. C. Example image of the SDS-PAGE of lysates of STX1-null neurons transduced with FLAG-tagged STX1A^WT^, STX1A^K260E^, or STX1A^CVCV^ loaded onto the SDS-PAGE after ABE method and visualized by HRP-Streptavidin antibody (top panel). After stripping the Streptavidin antibody, the membrane was developed with STX1A antibody (bottom panel). D. Example traces (left) and quantification of the amplitude (right) of EPSCs obtained from hippocampal autaptic STX1-null neurons rescued either with STX1A^WT^, STX1A^C271V^, STX1A^C272V^, or STX1A^CVCV^. E. Quantification of the decay time (80% - 20%) of the EPSC recorded from the same neurons as in (D). F. Example traces (left) and quantification of RRP recorded from the same neurons as in (D). G. Quantification of Pvr recorded from the same neurons as in (D). H. Example traces (left) and quantification of the frequency (right) of mEPSCs recorded from the same neurons as in (B). I. Example traces (left) and quantification (right) of STP measured by 50 stimulations at 10 Hz recorded from the same neurons as in (B). Data information: The artifacts are blanked in example traces in (D and F). The example traces in (H) were filtered at 1 kHz. In (D–H), data points represent single observations, the violin bars represent the distribution of the data with lines showing the median and the quartiles. In (I), data points represent mean ± SEM. Red annotations (stars) on the graphs show the significance comparisons to STX1A^WT^. Nonparametric Kruskal-Wallis test followed by Dunn’s *post hoc* test was applied to data in (D-H); *p ≤ 0.05, **p ≤ 0.01, ****p ≤ 0.0001. The numerical values are summarized in source data.

To test whether STX1A^K260E^ is palmitoylation deficient, we applied Acyl-Biotin-Exchange (ABE) method in which the palmitate group is exchanged by biotin through hydroxylamine (HAM) mediated cleavage of the thioester bond between a cysteine and a fatty acid chain (Brigidi and Bamji, 2013; Kang et al., 2008). The lysates of STX1-null neuronal cultures transduced with STX1A^WT^, STX1A^K260E^, STX1A^CVCV^ that are N-terminally tagged with FLAG epitope and non-transduced STX1-null neuronal cultures were treated with N-ethylmaleimide (NEM) solution to block the free thiol groups. STX1 was then pulled down using anti-FLAG magnetic beads. The beads that are attached to STX1 were then incubated in a solution either with or without HAM and subsequently with biotin solution. The covalently bound biotin to the free thiols now exposed after HAM cleavage of the thioester bonds between the palmitate and cysteine was detected by Western Blot using streptavidin antibody. A positive biotin band was detected only in STX1A^WT^ which was treated with the HAM solution (Figure 4C and Figure 4 - Supplement 1). Neither STX1A^K260E^ nor STX1A^CVCV^ produced any biotin positive band showing that both constructs lacked palmitoylation. We further probed the nitrocellulose membranes with STX1A antibody after stripping streptavidin antibody and observed that the lack of detection of biotinylated protein in the groups STX1A^K260E^ and STX1A^CVCV^ is not due to the loss of protein during the ABE-protocol (Figure 4C). Superimposition of the images acquired by streptavidin and STX1A antibody treatments showed that the biotin positive band in STX1A^WT^ lysates corresponds to STX1A (Figure 4 - Supplement 1).

We then analyzed electrophysiological properties of STX1A^C271V^ and STX1A^C272V^ that seemingly lack one palmitate group as well as STX1A^CVCV^ that lack both of its palmitate groups (Figure 4B). EPSCs recorded from STX1A^CVCV^ neurons trended towards a reduction from 6.60 nA to 4.58 nA on average with a p-value of 0.29 compared to that of STX1A^WT^. Neither STX1A^C271V^ nor STX1A^C272V^ showed a similar level of trend towards a reduction as they produced EPSCs of 5.16 nA with p-values of > 0.99 and of 0.83, respectively (Figure 4D). On the other hand, all the mutants trended towards having faster EPSC kinetics with a faster decay time with p-values ranging from 0.03 to 0.17 compared to that of STX1A^WT^ (Figure 4E). Together with an RRP size which is comparable among all the groups (Figure 4F), palmitoylation deficient neurons had significantly lower Pvr (Figure 4G), suggesting loss of palmitoylation impairs also the efficacy of Ca^2+^-evoked release. Consistent with a reduced Pvr, all palmitoylation deficient mutants showed almost no depression in the short-term plasticity (STP) as determined by 10 Hz stimulation (Figure 4I), suggesting an impairment in the vesicular release efficacy. Yet again, the most dramatic effect of palmitoylation deficiency was observed on spontaneous release as STX1A^C271V^ and STX1A^C272V^ mutants showed either a significant reduction in mEPSC frequency (STX1A^C272V^, p-value of 0.01) or a trend towards it (STX1A^C271V^, p-value of 0.15 (Figure 4H). Importantly, STX1A^CVCV^ mutant which lacks both palmitates showed almost no spontaneous neurotransmitter release (Figure 4H), a phenotype similar to STX1A^K260E^ mutant (Figure 2F).

Interestingly, the cysteine residues in STX1A’s TMD has been suggested to interact with presynaptic Ca^2+^-channels (Bachnoff et al., 2013; Cohen et al., 2007; Sajman et al., 2017; Sheng et al., 1994; Wiser et al., 1996) and to inhibit the baseline Ca^2+^-channel activity (Trus et al., 2001). This could underly the absence of mEPSC in STX1A^CVCV^ mutant as one mechanism proposed for spontaneous release is the stochastic opening of Ca^2+^-channels in the presynapse (Kaeser and Regehr, 2014; Williams and Smith, 2018). To test whether loss of palmitoylation by STX1A^K260E^ alone and loss of cysteine residues and their palmitoylation thereof would affect the Ca^2+^-channel activity, we monitored Ca^2+^-influx in the presynapse in the neurons additionally transduced with the Ca^2+^-sensor SynGCampf-6 by stimulating them with different numbers of APs (Figure 5A – 5C). As we have previously reported (Vardar et al., 2021), loss of STX1 reduced the global Ca^2+^-influx (Figure 5A and 5B). On the other hand, neither STX1A^CVCV^ nor STX1A^K260E^ showed significantly different SynGCampf-6 signal where the former trended towards an increase for 1AP and the latter trended towards a decrease for 2, 5, 10, and 20 APs (Figure 5A and 5C).

**Figure 5:**
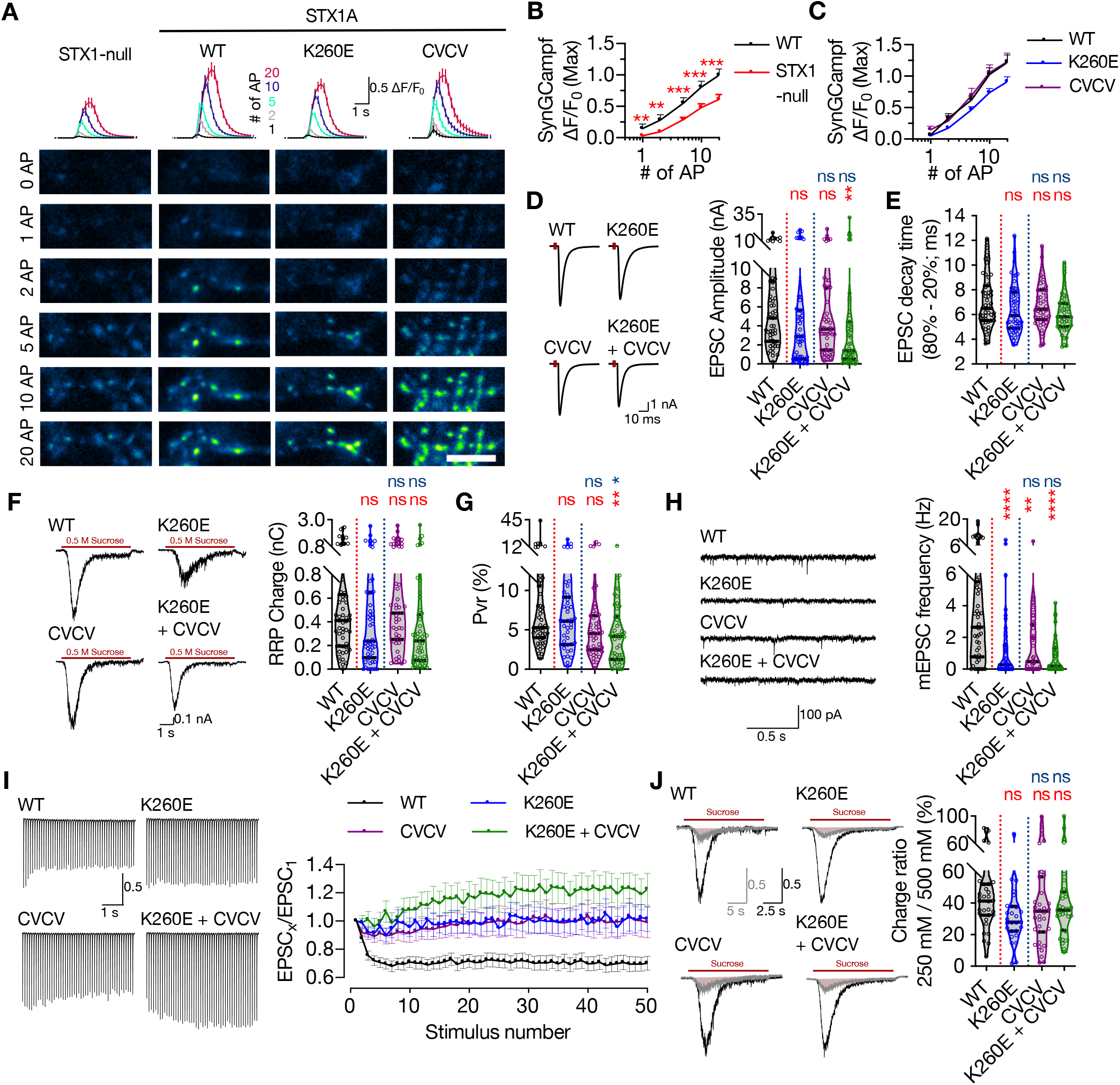
Combination of the K260E and CVCV mutations does not further change the phenotype of STX1A^K260E^. A. The average (top panel) of SynGCaMP6f fluorescence as (ΔF/F_0_) and example images thereof (bottom panels) in STX1-null neurons either not rescued or rescued with STX1A^WT^, STX1A^K260E^, or STX1A^CVCV^. The images were recorded at baseline, and at 1, 2, 5, 10, and 20 APs. Scale bar: 10 μm B. Quantification of the SynGCaMP6f fluorescence as (ΔF/F_0_) in STX1-null neurons either not rescued or rescued with STX1A^WT^. C. Quantification of the SynGCaMP6f fluorescence as (ΔF/F_0_) in STX1-null neurons rescued with STX1A^WT^, STX1A^K260E^, or STX1A^CVCV^. D. Example traces (left) and quantification of the amplitude (right) of EPSCs obtained from hippocampal autaptic STX1-null neurons rescued either with STX1A^WT^, STX1A^K260E^, STX1A^CVCV^, or STX1A^K260E+CVCV^. E. Quantification of the decay time (80% - 20%) of the EPSC recorded from the same neurons as in (D). F. Example traces (left) and quantification of RRP recorded from the same neurons as in (D). G. Quantification of Pvr recorded from the same neurons as in (D). H. Example traces (left) and quantification of the frequency (right) of mEPSCs recorded from the same neurons as in (D). I. Example traces (left) and quantification (right) of STP measured by 50 stimulations at 10 Hz recorded from same neurons as in (D). J. Example traces (left) and quantification (right) of the ratio of the charge transfer triggered by 250 mM sucrose over that of 500 mM sucrose recorded from same neurons as in (D) as a read-out of fusogenicity of the SVs. Data information: The artifacts are blanked in example traces in (D, F, and J). The example traces in (H) were filtered at 1 kHz. In (B, C, and I), data points represent mean ± SEM. In (D–H and J), data points represent single observations, the violin bars represent the distribution of the data with lines showing the median and the quartiles. Red annotations (stars) on the graphs show the significance comparisons to STX1A^WT^. Nonparametric Kruskal-Wallis test followed by Dunn’s *post hoc* test was applied to data in (B-H and J); **p ≤ 0.01, ***p ≤ 0.001, ****p ≤ 0.0001. The numerical values are summarized in source data.

So far, STX1A^K260E^ and STX1A^CVCV^ showed comparable phenotypes in synaptic neurotransmission as they both affected the spontaneous neurotransmitter release more drastically than the Ca^2+^-evoked release (Figure 2 and 4). However, only STX1A^K260E^ reduced the size of the RRP but not STX1A^CVCV^ (Figure 2 and 4). To uncouple the effects of the synaptic phenotype of K260E by means of the alterations on the membrane proximal electrostatic landscape by charge reversal mutation and by loss of palmitoylation, we created a STX1A construct in which K260E mutation was coupled with CVCV mutation (STX1A^K260E+CVCV^; Figure 4B). EPSC amplitudes recorded from STX1A^K260E+CVCV^ neurons were significantly smaller compared to that of STX1A^WT^ but not to that of STX1A^K260E^ (Figure 5D). Whereas the RRP size measured from STX1A^K260E+CVCV^ neurons was comparable to that of STX1A^K260E^, Pvr was significantly reduced (Figure 5F and 5G). Most importantly, spontaneous release was again dramatically reduced in STX1A^K260E+CVCV^ neurons (Figure 5H). Lastly, all the STX1A^K260E^, STX1A^CVCV^, and STX1A^K260E+CVCV^ mutants showed impairment in short-term plasticity as they showed either facilitation or almost no plasticity as opposed to the short-term depression observed in STX1A^WT^ neurons (Figure 5I). The observed alteration in the STP was not due to changes in the vesicle fusogenicity as proxied as the fraction of the RRP released by sub-saturating 250 mM sucrose solution (Figure 5J).

Because STX1A^K260E^ and STX1A^CVCV^ so far showed only a trend towards a reduction in EPSC amplitude, we pooled all the data obtained from STX1A^K260E^ (Figure 2, 5, and 6) and STX1A^CVCV^ (Figure 4 and 5) neurons and plotted values as normalized to STX1A^WT^ for each individual culture (Figure 5 - Supplement 1). EPSC amplitude was significantly reduced for both STX1A^K260E^ and STX1A^CVCV^ mutants in the pooled data (Figure 5 - Supplement 1).

**Figure 6:**
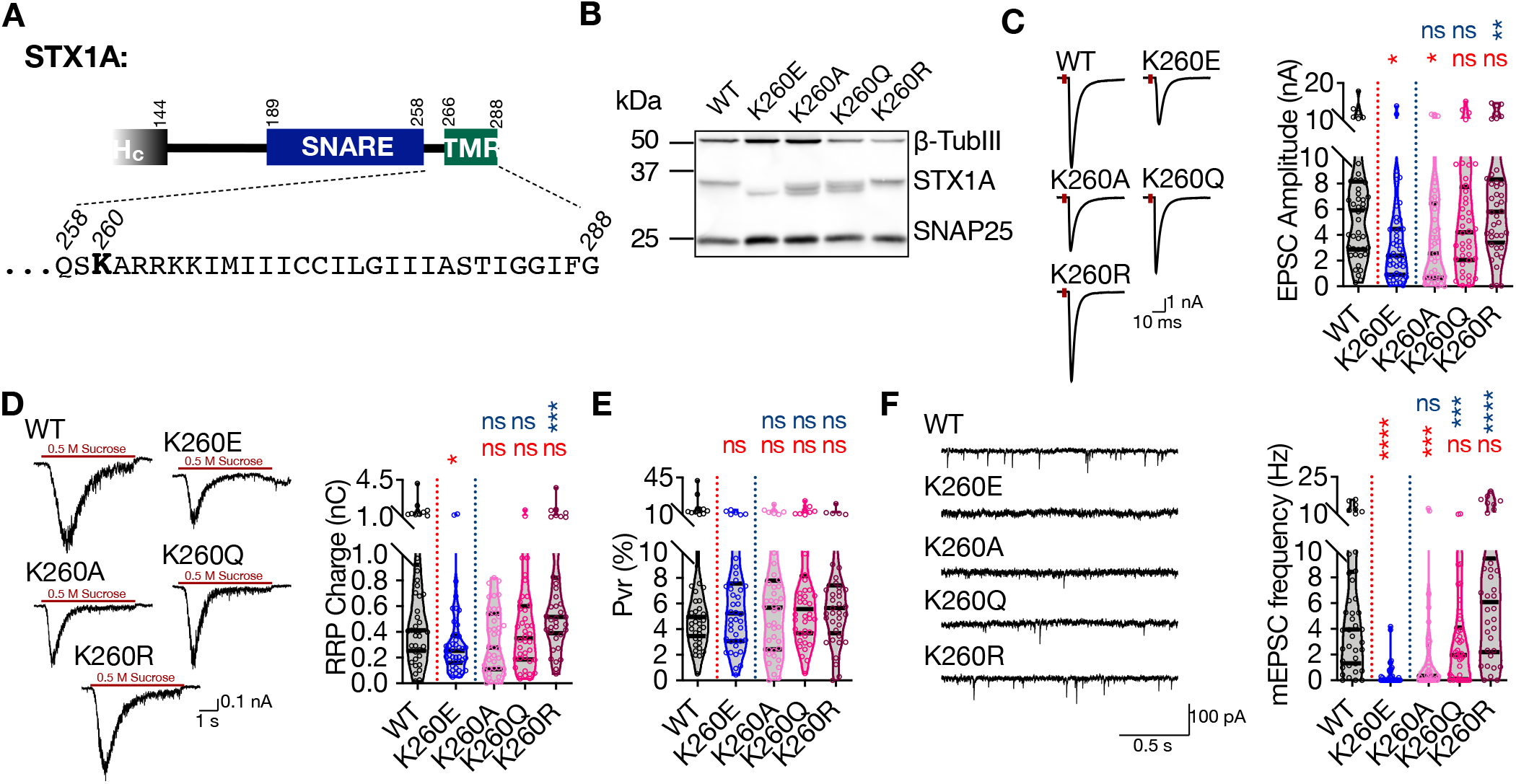
Palmitoylation of STX1A’s TMD depends on the presence of a basic residue at position aa 260 on its JMD. A. Position of the aa 260 on STX1A’s JMD. B. Example image of SDS-PAGE of the electrophoretic analysis of lysates obtained from STX1-null neurons transduced with STX1A^WT^, STX1A^K260E^, STX1A^K260A^, STX1A^K260Q^, or STX1A^K260R^. C. Example traces (left) and quantification of the amplitude (right) of EPSCs obtained from hippocampal autaptic STX1-null neurons rescued either with STX1A^WT^, STX1A^K260E^, STX1A^K260A^, STX1A^K260Q^, or STX1A^K260R^. D. Example traces (left) and quantification of RRP recorded from the same neurons as in (C). E. Quantification of Pvr recorded from the same neurons as in (C). F. Example traces (left) and quantification of the frequency (right) of mEPSCs recorded from the same neurons as in (C). Data information: The artifacts are blanked in example traces in (C and D). The example traces in (F) were filtered at 1 kHz. In (C–F), data points represent single observations, the violin bars represent the distribution of the data with lines showing the median and the quartiles. Red and blue annotations (stars and ns) on the graphs show the significance comparisons to STX1A^WT^ and STX1A^K260E^, respectively. Nonparametric Kruskal-Wallis test followed by Dunn’s *post hoc* test was applied to data in (C-F); *p ≤ 0.05, **p ≤ 0.01, ***p ≤ 0.001, ****p ≤ 0.0001. The numerical values are summarized in source data.

Next, we tested whether the palmitoylation deficiency is due to the loss of the lysine or due to the loss of a basic residue at position aa 260 (Figure 6A). For that purpose, we created STX1A mutants in which the lysine K260 on STX1A was exchanged either with a neutral and small alanine (K260A), a neutral glutamine (K260Q) which is more similar to glutamate, or with an arginine (K260R) which is basic. Whereas STX1A^K260A^ produced a banding pattern on SDS-PAGE with two lower bands (Figure 6B) similar to STX1A^R262E^ (Figure 3A), STX1A^K260Q^ produced three bands (Figure 6B) similar to STX1A^K264E^ (Figure 3A). On the other hand, STX1A^K260R^ was fully capable of rescuing the banding pattern of STX1A to the highest single band level similar to STX1A^WT^ (Figure 6B). Both EPSCs produced by STX1A^K260A^ and STX1A^K260Q^ did not significantly differ from those produced by STX1A^K260E^ (Figure 6C). On the other hand, STX1A^K260R^ significantly rescued the EPSC back to WT-like level (Figure 6C). Similarly, only STX1A^K260R^ neurons had significantly larger RRPs compared to the STX1A^K260E^ neurons (Figure 6D) and none of the mutants altered Pvr (Figure 6E). Remarkably, on the other hand, spontaneous release showed a graded improvement in the order of K260E < K260A < K260Q < K260R (Figure 6F) comparable to the banding pattern in SDS-PAGE (Figure 6B). This suggests that it is the presence of a basic residue at the position 260 in the JMD of STX1A that is important for palmitoylation of STX1A in its TMD and thereby for the regulation of spontaneous neurotransmitter release.

### The impacts of palmitoylation of STX1A’s TMD on spontaneous release and its regulation by STX1A’s JMD can be emulated by using different syntaxin isoforms

So far, we have shown that the basic nature of STX1A’s JMD plays an important and differential role in the regulation of the Ca^2+^-evoked and spontaneous release as well as vesicle priming (Figure 2 and 6) not only through its electrostatic interactions with the plasma membrane but also through its potential effects on the palmitoylation state of STX1A’s TMD. The JMD of STX1s is highly conserved across a number of species yet it shows variability among the different STX isoforms, which are involved in various intracellular trafficking pathways (Van Komen et al., 2005).

Among the members of the syntaxin family, only STX1s and STX3B, which share the same domain structure with a 67% sequence homology, have defined functions in presynaptic neurotransmitter release. Whereas STX1s are expressed in the central synapses that show ‘phasic’ release, STX3B is the predominant isoform in ‘tonically releasing’ retinal ribbon synapses where STX1 is excluded (Curtis et al., 2010; Curtis et al., 2008). Strikingly, however, STX3B has a glutamate (E) in its JMD at the position 259 leading to the sequence “EARRKK” (Fig. 7A). It is striking that this is the sequence we found which renders STX1A incompetent for spontaneous release potentially through its incompatibility for the palmitoylation of STX1A’s TMD (Figure 2, 5, and 6). To test how naturally having a glutamate residue as its most N-terminal residue of its JMD effects the neurotransmitter release properties of STX3B in a central synapse, we expressed STX3B in STX1-null hippocampal neurons with or without the charge reversal mutation E259K, which produces a STX1A-like JMD in STX3B (Figure 7A).

**Figure 7:**
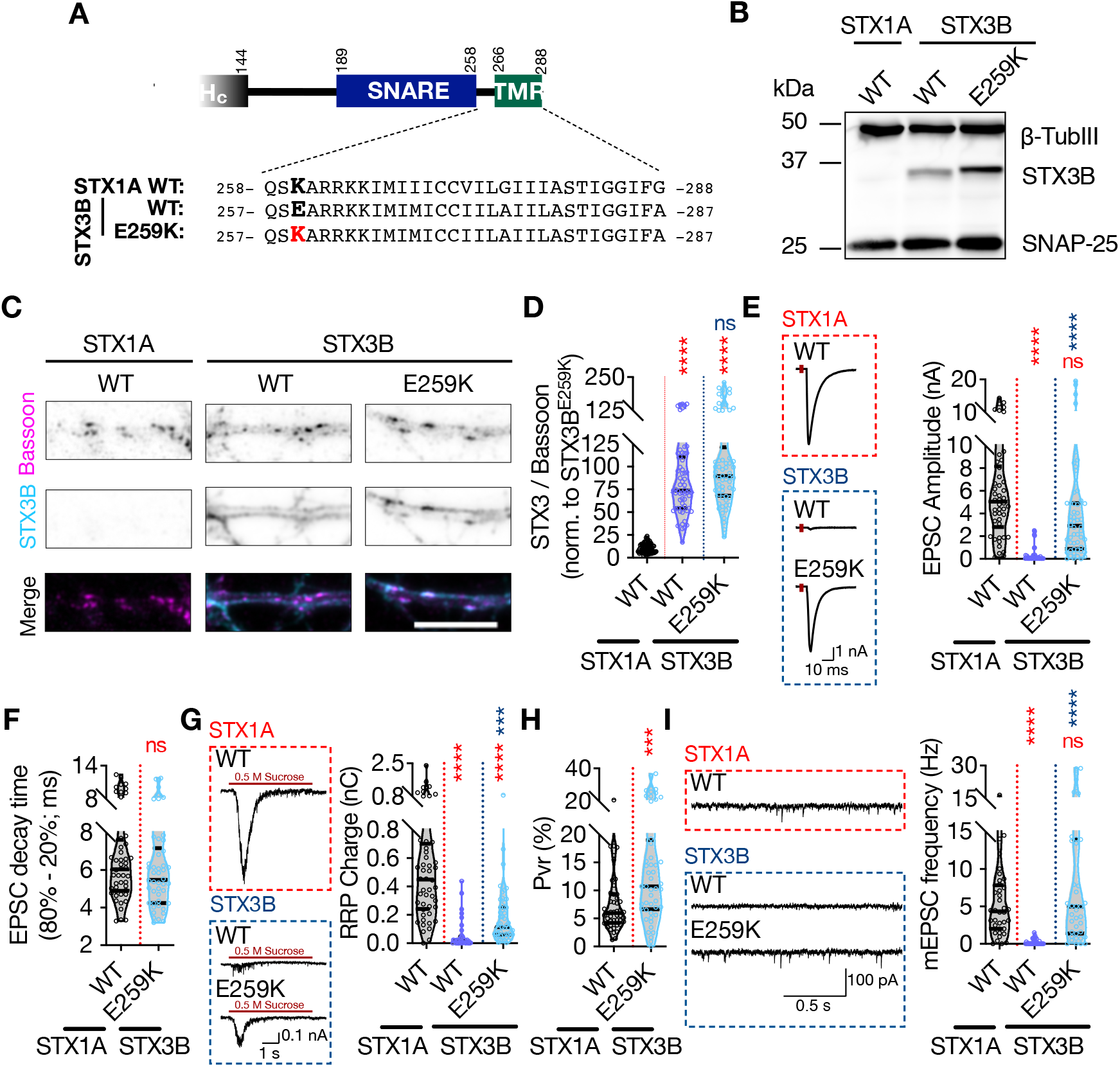
Retinal ribbon specific STX3B has a glutamate at position aa 259 rendering its JMD similar to that of STX1AK260E and E259K mutation on STX3B acts as a molecular on-switch. A. Position of the aa 259 on STX3B’s JMD. B. Example image of SDS-PAGE of the electrophoretic analysis of lysates obtained from STX1-null neurons transduced with STX1A^WT^, STX3B^WT^, or STX3B^E259K^. C. Example images of immunofluorescence labeling for Bassoon and STX3B, shown as magenta and cyan, respectively, in the corresponding composite pseudocolored images obtained from high-density cultures of STX1-null hippocampal neurons rescued with STX1A^WT^, STX3B^WT^, or STX3B^E259K^. Scale bar: 10 μm D. Quantification of the immunofluorescence intensity of STX3B as normalized to the immunofluorescence intensity of Bassoon in the same ROIs as shown in (C). The values were then normalized to the values obtained from STX3B^WT^ neurons. E. Example traces (left) and quantification of the amplitude (right) of EPSCs obtained from hippocampal autaptic STX1-null neurons rescued with STX1A^WT^, STX3B^WT^, or STX3B^E259K^. F. Quantification of the decay time (80% - 20%) of the EPSC recorded from the same neurons as in (E). G. Example traces (left) and quantification of RRP recorded from the same neurons as in (E). H. Quantification of Pvr recorded from the same neurons as in (E). I. Example traces (left) and quantification of the frequency (right) of mEPSCs recorded from the same neurons as in (E). Data information: The artifacts are blanked in example traces in (E and G). The example traces in (I) were filtered at 1 kHz. In (D–I), data points represent single observations, the violin bars represent the distribution of the data with lines showing the median and the quartiles. Red and blue annotations (stars and ns) on the graphs show the significance comparisons to STX1A^WT^ and STX3B^WT^, respectively. Nonparametric Kruskal-Wallis test followed by Dunn’s *post hoc* test was applied to data in (C-F); ***p ≤ 0.001, ****p ≤ 0.0001. The numerical values are summarized in source data.

Firstly, we probed STX3B^WT^ and STX3B^E259K^ on SDS-PAGE to test whether the E259K mutation changes the banding pattern of STX3B and observed that STX3B^E259K^ indeed showed a higher molecular weight compared to that of STX3B^WT^ (Figure 7B). This is consistent with the expected palmitoylation deficiency of STX3B^WT^ due to the presence of a glutamate at the position which corresponds to K260 in STX1A. Before proceeding with our electrophysiological recordings, we also assessed the expression level of STX3B^E259K^ relative to the expression of STX3B^WT^ and determined that both constructs are exogenously expressed at comparable levels at Bassoon-positive puncta when lentivirally introduced into STX1-null neurons (Figure 7C and 7D). Endogenous expression of STX3B in STX1A^WT^ neurons did not produce a measurable signal at the exposure times of the excitation wavelength used in this study (Figure 7C and 7D).

Next, we tested how the replacement of STX1s either with STX3B^WT^ or STX3B^E259K^ affects the neurotransmitter release by measuring the Ca^2+^-evoked and spontaneous release and the RRP of the SVs (Figure 7E – 7I). STX3B^WT^ was unable to rescue neither form of neurotransmitter release nor vesicle priming in STX1-null neurons deeming it dysfunctional in conventional synapses even though it efficiently mediates the neurotransmitter release from retinal ribbon synapses (Figure 7E – 7I). Surprisingly, however, E259K mutation served as a molecular on-switch for STX3B as STX3B^E259K^ fully rescued both Ca^2+^-evoked release with normal release kinetics and spontaneous release (Figure 7E, 7F, and 7I). On the other hand, the size of the RRP in STX3B^E259K^ neurons remained at ~50% of that observed in STX1A^WT^ neurons (Figure 7G), which then led to an increased Pvr (Figure 7H). This suggests that the N-terminal lysine of the JMD indeed plays a vital role in the functioning of neuronal syntaxins.

The retinal ribbon synapse specific STX3B is a splice variant of STX3A which is also a neuronal syntaxin with roles indicated in postsynaptic exocytosis (Jurado et al., 2013). The differential splicing of STX3A and STX3B occurs in the middle of the SNARE domain generating two products that are identical at their regions between the layer 0 of the SNARE domain and the N-terminus of the protein (Figure 8 - Supplement 1). On the other hand, the rest of these proteins spanning the C-terminal half of their SNARE domains, JMD, and TMD show only a 43.1% homology (Figure 8A and Figure 8 - Supplement 1). Among the sequence differences between the STX3B and STX3A, the JMD and the TMD are of great importance as STX3A lacks the cysteine residues in its TMD and thus the substrate for palmitoylation. Furthermore, its JMD not only has a glutamine at the region corresponding to K260 in STX1A but also has one less basic residue compared to that of STX1A and STX3B (Figure 8A and Figure 8 - Supplement 1).

**Figure 8:**
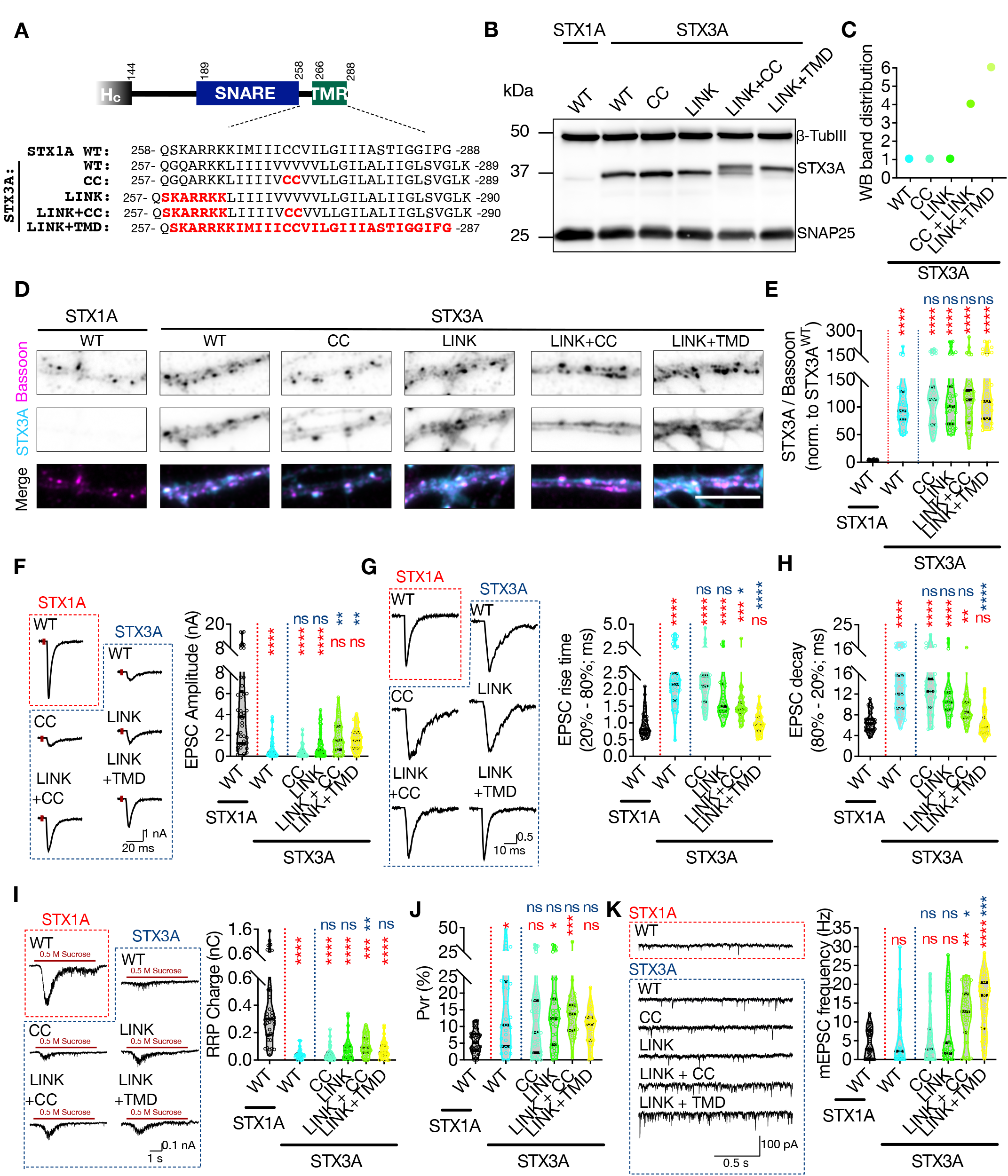
The impacts of palitoylation of STX1A’s TMD on spontaneous release and can be emulated by using STX3A. A. Comparison of the JMD and TMD regions of STX1A and STX3A and the mutations introduced into STX3A. B. Example image of SDS-PAGE of the electrophoretic analysis of lysates obtained from STX1-null neurons transduced with STX1A^WT^, STX3A^WT^, STX3A^CC^, STX3A^LINK+CC^, or STX3A^LINK+TMR^. C. Quantification of the STX3A band pattern on SDS-PAGE of neuronal lysates through assignment of arbitrary hierarchical numbers from 1 to 6 based on the distance traveled as in Figure 3A. D. Example images of immunofluorescence labeling for Bassoon and STX3A, shown as magenta and cyan, respectively, in the corresponding composite pseudocolored images obtained from high-density cultures of STX1-null hippocampal neurons rescued with STX1A^WT^, STX3A^WT^, STX3A^CC^, STX3A^LINK+CC^, or STX3A^LINK+TMR^. Scale bar: 10 μm E. Quantification of the immunofluorescence intensity of STX3A as normalized to the immunofluorescence intensity of Bassoon in the same ROIs as shown in (D). The values were then normalized to the values obtained from STX3A^WT^ neurons. F. Example traces (left) and quantification of the amplitude (right) of EPSCs obtained from hippocampal autaptic STX1-null neurons rescued with STX1A^WT^, STX3A^WT^, STX3A^CC^, STX3A^LINK+CC^, or STX3A^LINK+TMR^. G. Example traces with the peak normalized to 1 (left) and quantification of the EPSC rise time measured from 20% to the 80% of the EPSC recorded from STX1-null neurons as in (F). H. Quantification of the decay time (80% - 20%) of the EPSC recorded from the same neurons as in (F). I. Example traces (left) and quantification of RRP recorded from the same neurons as in (F). J. Quantification of Pvr recorded from the same neurons as in (F). K. Example traces (left) and quantification of the frequency (right) of mEPSCs recorded from the same neurons as in (F). Data information: The artifacts are blanked in example traces in (F, G, and I). The example traces in (K) were filtered at 1 kHz. In (E–K), data points represent single observations, the violin bars represent the distribution of the data with lines showing the median and the quartiles. Red and blue annotations (stars and ns) on the graphs show the significance comparisons to STX1A^WT^ and STX3A^WT^, respectively. Nonparametric Kruskal-Wallis test followed by Dunn’s *post hoc* test was applied to data in (C-F); *p ≤ 0.05, **p ≤ 0.01, ***p ≤ 0.001, ****p ≤ 0.0001. The numerical values are summarized in source data.

To test the effects of the palmitoylation of the TMD and the basic residues in the JMD regardless of the SNARE domain, we created STX3A mutants in which either two cysteine residues were incorporated into its TMD (STX3A^CC^) or in which the ‘KARRKK’ sequence was introduced into its JMD (STX3A^LINK^) to transmute this region of STX3A into STX1A-like (Figure 8A). Furthermore, to evaluate the interplay between the JMD and the palmitoylation of STXs, we also created two additional mutants in which cysteine and JMD incorporation was combined (STX3A^LINK + CC^) or in which the whole region spanning the JMD and the TMD of STX3A was exchanged with the corresponding region of STX1A (STX3A^LINK + TMD^) (Figure 8A).

We again probed the STX3A^WT^ and the STX3A mutants on SDS-PAGE to test for possible alterations in the banding pattern and thereby the palmitoylation state of STX3A (Figure 7B). Consistent with our proposal that there is an interplay between the JMD of STX1A and its palmitoylation in its TMD, we observed STX3A^WT^ produced one single band and that was not affected when the CC or STX1A’s JMD alone were incorporated in STX3A (Figure 8B and 8C). However, combination of CC and STX1A’s JMD in STX3A was enough to reach some partial rescue of the palmitoylation state of STX3A, whereas the introduction of the whole region of STX1A spanning its JMD and TMD effectively reached the full palmitoylation state as manifested by the production of higher molecular weight bands on SDS-PAGE (Figure 8B and 8C). Whereas we could not detect endogenous expression of STX3A in STX1A^WT^ neurons at the exposure times used for the excitation wavelength tested, none of the STX3A mutations altered its exogenous expression level at Bassoon-positive puncta compared to that of STX3A^WT^ (Figure 8D and 8E).

Remarkably, STX3A^WT^, albeit not being a member of any presynaptic neurotransmitter release machinery, supported some level of neurotransmitter release when lentivirally expressed in STX1-null neurons (Figure 8F), unlike STX3B^WT^ (Figure 7E). Whereas the EPSC amplitudes recorded from STX3A^WT^ neurons reached ~15% of that of STX1A^WT^ neurons (Figure 8F), the EPSCs were significantly slowed down as shown by a doubled duration of both the EPSC rise and the EPSC decay compared to those recorded from STX1A^WT^ neurons (Figure 8G and 8H). Introduction of the two cysteine residues or the JMD into STX3A did not lead to any enhancement in its efficacy to support neurotransmitter release, as both the STX3A^CC^ and STX3A^LINK^ neurons produced EPSCs comparable to STX3A^WT^ both in size and kinetics (Figure 8F – 8H). Interestingly, on the other hand, both STX3A^LINK+CC^ and STX3A^LINK+TMD^ generated EPSCs which were significantly bigger compared to that recorded from STX3A^WT^ and which showed only a trend towards a reduction compared to that of produced by STX1A^WT^ (Figure 8F). Remarkably, the synchronicity of EPSC was rescued partially by STX3A^LINK+CC^ and fully by STX3A^LINK+TMD^ (Figure 8G and 8H) pointing out the involvement of not only STX1A’s TMD in the synchronicity of Ca^2+^-evoked release but also of its JMD and its palmitoylation. Moreover, neurons in which STX1s were exogenously replaced by STX3A^WT^ harbored only a very small RRP and that was not rescued by any STX3A mutant (Figure 8I) leading to a high Pvr in all neurons expressing STX3A constructs (Figure 8J).

Surprisingly, STX3A^WT^ neurons spontaneously released SVs at a frequency comparable to that of STX1A^WT^ neurons and introduction of the CC or the LINK mutations alone into STX3A showed no alterations in the mEPSC frequency (Figure 8K). Remarkably, combination of the linker region mutation either with CC mutation or with the whole TMD and thereby allowing the palmitoylation of the cysteines in the TMD spontaneously discharged the SVs at a frequency that reached 3-4-fold of that of spontaneous release from STX1A^WT^ neurons (Figure 8K). This substantial increase in the mEPSC frequency driven by the palmitoylation of the STX3A’s TMD essentially points out the same mechanism for the regulation of the spontaneous release by the palmitoylation of STX1A’s TMD, as its inhibition either by STX1A^K260E^ or STX1A^CVCV^ diminished spontaneous release (Figure 2, 4, 5, and 6). This suggests that the prevention of the palmitoylation of the CC residues in the TMD of syntaxins blocks the spontaneous release.

As mentioned above, retinal ribbon synaptic STX3B and postsynaptic STX3A are splice variants which differ only at their C-terminus from the layer 0 of the SNARE domain to the end of the TMD (Figure 8 - Supplement 1). The TMD of STX3B is more STX1A-like as the comparative sequence alignment shows that all the amino acids at respective positions in the TMD region of STX3B and STX1A share similar biophysical properties if not the same (Figure 8 - Supplement 1). However, the STX3B^E259K^, which can be considered as the active form of STX3B in conventional synapses, did not show higher spontaneous release efficacy compared to STX1A, even though the cysteine residues are present in that mutant (Figure 7). On the other hand, STX3A^LINK+TMD^, which has STX1A’s TMD that is now more similar to STX3B, shows an unclamped spontaneous release of SVs (Figure 8H). Taken these into account, we compared the amino acid sequences of STX1A, STX3B^E259K^, and STX3A^LINK+TMD^ and plotted the mEPSC frequencies as values normalized to the STX1A^WT^ for each individual culture (Figure 8 - Supplement 1). The sequence alignment shows that the STX3A^LINK+TMD^ mutant which spontaneously releases SVs at the highest frequency differs from both STX1A^WT^ and STX3B^E259K^ only in the C-terminal half of its SNARE domain (Figure 8 - Supplement 1). This suggests also the C-terminal half of STX1A’s SNARE domain from its layer 0 to layer 8 plays a pivotal role in clamping of the spontaneous vesicle fusion.

A line of hypotheses in account of spontaneous vesicle fusion suggests that the vesicles close to the Ca^2+^-channels fuse with the membrane upon stochastic opening of Ca^2+^-channels (Kaeser and Regehr, 2014; Williams and Smith, 2018). Therefore, alterations in the regulation of Ca^2+^-channel gating, such as proposed inhibition of baseline activity of Ca^2+^-channels by the cysteine residues of STX1A’s TMD (Trus et al., 2001), could lead to a decrease or increase in the spontaneous neurotransmitter release as a result of altered spontaneous Ca^2+^-influx into the synapse. On the other hand, desynchronization of the Ca^2+^-evoked neurotransmitter release by STX3A^WT^ might be a result of uncoupling of the Ca^2+^-channel-SV spatial organization. In that scenario, it is plausible that the TMD of STX1A plays a role in the Ca^2+^-channel-SV coupling, as its incorporation into STX3A leads to the recovery of synchronous release (Figure 8G and 8H). To test whether and how STX3B and STX3A affect the global Ca^2+^-influx in their WT or mutant forms, we again co-expressed the Ca^2+^-sensor SynGCampf6 in the autaptic synapses and measured the Ca^2+^-influx upon differing numbers of APs (Figure 9). Interestingly, both STX3B^WT^ and STX3A^WT^ failed to rescue the STX1-loss driven reduction of the Ca^2+^-influx in the presynaptic terminals (Figure 9A and 9B). On the other hand, STX3B^E259K^ rescued the global Ca^2+^-influx at higher number of APs elicited, suggesting that syntaxins indeed might play a role in the gating of Ca^2+^-channels (Figure 9A). Surprisingly, however, STX3A mutants did not have any impact on Ca^2+^-influx as it remained at STX1-null levels even for the mutants that rescue the EPSC synchronicity and that lead to excessive amount of spontaneous neurotransmitter release (Figure 9B). As the Ca^2+^-influx in neurons expressing STX3A variants remained low at any AP number compared to that of STX1A neurons, it is likely that the overall Ca^2+^-trafficking and hence Ca^2+^-abundance might be impaired in the absence of STX1 which cannot be rescued by STX3A.

**Figure 9:**
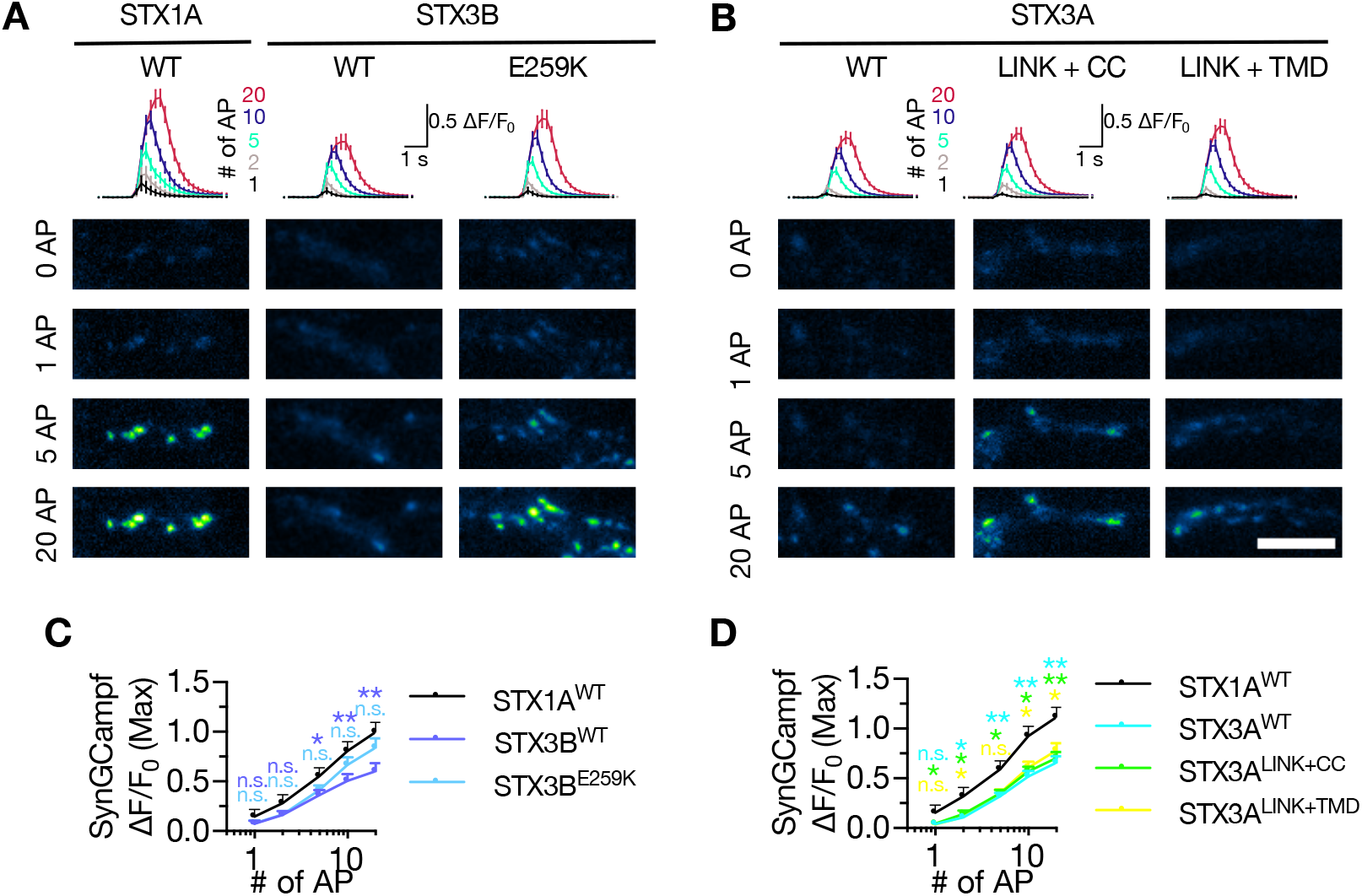
Neither STX3A nor STX3B rescues the global Ca2+-influx back at WT-like level in STX1-null neurons. A. The average (top panel) of SynGCaMP6f fluorescence as (ΔF/F_0_) and example images thereof (bottom panels) in STX1-null neurons rescued with STX1A^WT^, STX3B^WT^, or STX3B^E259K^. The images were recorded at baseline, and at 1, 2, 5, 10, and 20 APs. Scale bar: 10 μm B. The average (top panel) of SynGCaMP6f fluorescence as (ΔF/F_0_) and example images thereof (bottom panels) in STX1-null neurons rescued with STX1A^WT^, STX3A^WT^, STX3A^CC^, STX3A^LINK+CC^, or STX3A^LINK+TMR^. The images were recorded at baseline, and at 1, 2, 5, 10, and 20 APs. Scale bar: 10 μm C. Quantification of the SynGCaMP6f fluorescence as (ΔF/F_0_) in STX1-null neurons rescued with STX1A^WT^, STX3B^WT^, or STX3B^E259K^. D. Quantification of the SynGCaMP6f fluorescence as (ΔF/F_0_) in STX1-null neurons rescued with STX1A^WT^, STX3A^WT^, STX3A^CC^, STX3A^LINK+CC^, or STX3A^LINK+TMR^. Data information: In (C and D), data points represent mean ± SEM. All annotations (stars and ns) on the graphs show the significance comparisons to STX1A^WT^ with the color of corresponding group. Nonparametric Kruskal-Wallis test followed by Dunn’s *post hoc* test was applied to data in (C and D); *p ≤ 0.05, **p ≤ 0.01. The numerical values are summarized in source data.

## Discussion

The presynaptic SNARE complex formation by STX1A, SYB2, and SNAP-25 unequivocally set up the vesicular and plasma membrane in close proximity through the N-to-C zippering of their SNARE domains (Gao et al., 2012; Sorensen et al., 2006; Stein et al., 2009). In our study we addressed whether STX1A play additional roles in vesicle fusion to facilitate the membrane merger through its JMD and TMD. Based on our data we propose that the JMD of STX1A indeed regulates the membrane merger through adjusting the distance and the electrostatic nature of the inter-membrane area along the *trans*-SNARE complex. Whereas STX1A’s JMD also determines the palmitoylation state of STX1A’s TMD, the palmitoylation of STX1A’s TMD might directly influence the energy barrier for membrane merger, which is specifically apparent for spontaneous release. We conclude that the JMD and TMD of STX1A do not have a function merely as a membrane anchor but they are actively involved in the regulation of vesicle fusion.

### The role of STX1A’s JMD in neurotransmitter release extends beyond its interaction with PIP2

The current models of SV fusion have in fact given a significant role to the JMD of STX1A in vesicular fusion, where it drives STX1A clustering through its interaction with PIP2/PIP3 (Khuong et al., 2013; van den Bogaart et al., 2011a) and thereby indirectly regulates the spatial organization of vesicle docking as PIP2 also binds to SYT1 (Aoyagi et al., 2005; Chen et al., 2021; Honigmann et al., 2013; Park et al., 2015). However, the C-terminal lysine residues in STX1A’s JMD play a more prominent role in PIP2 binding than the intermediate arginine residues do (Murray and Tamm, 2011) and neutralization of both K264 and K265 leads to a significant reduction in neurotransmitter release in fly neurons (Khuong et al., 2013). This is not the case for our charge reversal STX1A mutants as the C-terminal lysine to glutamate mutations in STX1A^K264E^ and STX1A^K265E^ that are assumed to have the greatest impact on PIP2 binding (Murray and Tamm, 2011) showed the mildest phenotype in neurotransmitter release (Figure 2). Furthermore, the other charge reversal mutants, STX1A^R262E^, STX1A^R263E^, and STX1A^K260E^ showed differential effects on spontaneous and Ca^2+^-evoked release (Figure 2), which is not consistent with the hypothesis of a general reduction of vesicle docking and priming due to a perturbed STX1A-PIP2/PIP3 interaction but with a general impairment in the electrostatic regulation of the membrane merger. Importantly, the sensitivity of SYB2’s JMD to the charge neutralization mutations (Williams et al., 2009) further corroborates this notion.

Remarkably, our data also show that the function of STX1A’s JMD is not limited to the regulation of the electrostatic nature of the intermembrane area, but also possibly to the coordination of the intermembrane distance along the *trans*-SNARE complex. The slowing down of the release together with the unclamping of spontaneous release by STX1A^GSG265^ (Figure 1) is intriguingly reminiscent of the phenotype of the loss of the Ca^2+^-sensor SYT1 (Chang et al., 2018; Courtney et al., 2019; Vevea and Chapman, 2020; Xu et al., 2009). Through its interactions with the lipids, SYT1 primarily assists the closure of the gap between the two membranes for tight docking of the SVs (Arac et al., 2006; Chang et al., 2018; Chen et al., 2021; van den Bogaart et al., 2011b) and thereby accelerates the Ca^2+^-evoked fast synchronous release (Geppert et al., 1994; Littleton et al., 1993). Thus, it is possible that an increase in the intermembrane distance along the *trans*-SNARE complex by an elongation of the STX1A’s JMD even by one helical turn and thus less than 1 nm might impair SYT1’s function as a distance regulator in synchronizing the vesicular release.

It is not clear, however, why STX1A^GSG259^ has a more deleterious effect on Ca^2+^-evoked release than STX1A^GSG265^ does, but it is conceivable that it blocks the helical continuity of the SNARE complex into the JMD of STX1A and SYB2. Remarkably, this phenomenon of asymmetrical impact of SNARE-JMD or JMD-TMD decoupling on vesicle fusion is not unique to STX1A as it is also observed by SYB2 mutants in that the JMD was also elongated at different positions (Hu et al., 2020; Kesavan et al., 2007; McNew et al., 1999; Mostafavi et al., 2017). Similarly, helix breaking proline insertion at the junction of SNARE-JMD but not at the junction of JMD-TMD of SYB2 is detrimental to liposome fusion (Hu et al., 2020), suggesting an essential mechanistic similarity between the regulation of the membrane merger by the JMD of the plasma and the vesicular SNAREs. Given that the spontaneous release was either unaffected or facilitated by the elongation of STX1A’s JMD (Figure 1), it appears that the JMD of both STX1A and SYB2 contribute to the vesicle fusion possibly by regulating the electrostatic nature and the distance of the intermembrane area along the *trans*-SNARE complex and the helical continuity of the SNARE complex into the SNARE JMD in a Ca^2+^-dependent manner.

### Palmitoylation of STX1A’s TMD potentially reduces the energy barrier required for membrane merger

Palmitoylation as a PTM is used for various functions of a wide panel of pre- and postsynaptic proteins (Kang et al., 2008; Matt et al., 2019; Naumenko and Ponimaskin, 2018; Prescott et al., 2009). Firstly, it anchors the soluble proteins, such as SNAP-25 and CSPα, into their target membranes (Prescott et al., 2009), which is generally considered as the primary function of palmitoylation. Palmitoylation of the integral membrane proteins such as STX1A, SYB2, and SYT1, however, has less defined functions (Prescott et al., 2009). It is known that palmitoylation dependent alteration of a TMD’s length can affect its hydrophobic mismatch status with the corresponding carrier membrane (Greaves and Chamberlain, 2007) and therefore it can be utilized for the hydrophobic mismatch regulated transport of an integral membrane protein through Golgi-complex (Ernst et al., 2018; Ernst et al., 2019). Based on the same mechanism, palmitoylation can also determine the final localization and the clustering of an integral membrane protein particularly in cholesterol rich, thick, and rigid membranes, which does not conform to the general properties of TMDs (Levental et al., 2010; Melkonian et al., 1999; van Duyl et al., 2002). In fact, STX1A forms clusters in the membrane that depend on its cholesterol content (Lang et al.; Murray and Tamm, 2009) besides its PIP2 content (Honigmann et al., 2013; Murray and Tamm, 2009). Furthermore, SYB2, which is also palmitoylated in its TMD (Kang et al., 2004; Veit et al., 2000), is associated with cholesterol-rich lipid rafts derived from SVs (Chamberlain et al., 2001; Chamberlain and Gould, 2002), which affects its SNARE domain and TMD conformation (Han et al., 2016; Tong et al., 2009; Wang et al., 2020).

Taken together with the reported reduction of Ca^2+^-evoked neurotransmitter release upon cholesterol depletion (Lang et al., 2001; Wasser et al.), it is tempting to speculate that the impacts of palmitoylation deficiency of STX1A’s TMD might stem from a general localization, oligomerization and mobility defect of STX1A, similar to the cause of neurotransmitter release deficiency in neurons expressing non-palmitoylated SYT1 (Kang et al., 2004). However, such a localization and mobility defect of STX1A would be detrimental not only to spontaneous neurotransmitter release but to all types of vesicle fusion and even vesicle priming. Yet, our palmitoylation deficient STX1A mutants do not manifest a drastic change in vesicle priming and Ca^2+^-evoked vesicle fusion (Figure 4) and chemical removal of cholesterol has been shown to increase spontaneous vesicle fusion (Wasser et al., 2007). These render the hypothesis of mislocalized STX1A due to the loss of palmitoylation unlikely. As loss of complete or individual palmitoylation sites on SNAP25 does not compromise its proper localization (Greaves et al., 2009; Vogel et al., 2000; Washbourne et al., 2001) but vesicle exocytosis (Washbourne et al., 2001), it can be argued that the palmitoylation of the t-SNARE proteins has a direct influence on the membrane merger.

Membrane merger is an energetically costly process and in central synapses several mechanisms are employed to lower the energy barrier for the SV fusion. These mechanisms include - besides the SNARE complex formation - the concerted actions of SYT1 and complexin (CPX), which confer the central synapse with a spatial and temporal acuity of SV release upon an incoming Ca^2+^-signal (Brunger et al., 2018; Rizo, 2018). Furthermore, the TMD of SYB2 itself also can facilitate the membrane merger through its tilted conformation with a high flexibility in the membrane that largely stems from its distinctively high propensity for β-branches (Dhara et al., 2016; Hastoy et al., 2017). Whether or not STX1A’s TMD, too, shows a higher propensity for β-sheets than for α-helices is not known, but likely, as it is rich in β-branching isoleucine (Figure 1). Remarkably, on the other hand, a similar function in TMD tilting has been also attributed to the palmitoylation of membrane proteins (Blaskovic et al., 2013; Charollais and Van Der Goot, 2009). In that light, we propose that the palmitoylation of STX1A’s TMD serves as another mechanism for the lowering of the energy barrier for membrane fusion. This mechanism might seem particularly important for spontaneous vesicle fusion at first glance. However, the reduction of also the Ca^2+^-evoked release albeit being small emphasizes a general impairment in the vesicle fusion due to loss of palmitoylation. This is more evident in the decrease in the vesicular release probability especially revealed by the impairment in STP elicited by a 10 Hz stimulation in neurons that express STX1A single or double palmitoylation mutants as well as STX1A^K260E^ (Figure 4 and 5).

Another support for our hypothesis that palmitoylation of STX1A’s TMD potentially reduces the energy barrier for membrane merger comes from our observation that the incorporation of the palmitoylation substrate cysteines into the TMD of STX3A exclusively facilitates the spontaneous fusion even when the rest of STX3A’s TMD is kept unchanged (Figure 8). According to one line of evidence, spontaneous and Ca^2+^-evoked vesicle fusion share the same mechanism and membrane topology and thus the stochastic opening of Ca^2+^-channels enables the spontaneous fusion of single SVs that are in ‘fusion-ready’ state (Kaeser and Regehr, 2014; Williams and Smith, 2018). Taking this hypothesis into account, it can be proposed that natural palmitoylation of STX1A’s TMD and forced palmitoylation of STX3A’s TMD might increase the number of SVs that more easily overcome the energy barrier for membrane merger. Alternatively, it is also proposed that spontaneous and Ca^2+^-evoked release are mechanistically and spatially distinguished (Guzikowski and Kavalali, 2021; Kaeser and Regehr, 2014; Kavalali, 2015) and thus it can be suggested that palmitoylation of STX1A’s TMD drives it to the release sites allocated for spontaneous vesicle fusion. This implies a dynamic regulation of STX1A palmitoylation based on needs of spontaneous release. However, the turn-over of STX1A palmitoylation seems to be slow (Kang et al., 2008), similar to the turn-over of SNAP25 and SYT1 palmitoylation (Heindel et al., 2003; Kang et al., 2004; Kang et al., 2008), which is not compatible with the scenario of a dynamic and spatial regulation of spontaneous release through differential localization of STX1A based on its palmitoylation state.

### STX1A’s TMD is required for the synchronicity of the Ca^2+^-evoked release

Our studies using the charge reversal mutations on the JMD and the palmitoylation mutations on the TMD of STX1A show that these mutants can reach the AZ, as all of them could mediate neurotransmitter release of some sort (Figure 2, 4, 5, and 6). However, a potential effect of STX1A’s TMD for a fine tuning of its localization is not completely beyond comprehension, as the differential distribution of other syntaxins, STX3A and STX4, in the plasma membrane is based on their TMD length (Bulbarelli et al., 2002; Watson and Pessin, 2001). Consistently, swapping STX3A’s TMD with STX1A’s TMD was sufficient to rescue the synchronous release (Figure 8) suggesting a role for STX1A’s TMD in proper Ca^2+^-secretion-vesicle fusion coupling. The TMD of the retinal ribbon synapse syntaxin, STX3B, on the other hand, is almost identical to STX1A’s TMD with only two amino acid difference in the sequence (Figure 8 – Supplementary Figure 1), further pointing out the importance of the TMD for syntaxins involved in neurotransmitter release either from conventional or ribbon synapses.

Surprisingly, STX3B’s JMD differ from that of STX1A in a way that it resembles to the mutant STX1A^K260E^ and the STX3B^E259K^ mutation acts as a molecular on-switch in the STX1-null hippocampal synapses (Figure 7). Why the alteration of the STX3B’s but not STX1A’s JMD has such a dramatic effect also on Ca^2+^-evoked vesicle fusion and vesicle priming is not clear. Yet, different syntaxins might go through different conformational changes due to different intramolecular interactions with their regulatory N-terminal domains. Indeed, STX3B’s opening in retinal ribbon synapses, where Munc13s do not function as a primary vesicle priming factor (Cooper et al., 2012), is mediated through its N-peptide phosphorylation (Campbell et al., 2020; Liu et al., 2014). Whereas such an opening mechanism has not been reported for STX1A, the JMD has been proposed to induce conformational changes on STX1A through interactions with PIP2 which in turn leads to the phosphorylation of its N-peptide on S14 (Khelashvili et al., 2012; Singer-Lahat et al., 2018). Therefore, it is conceivable that E259K mutation’s impact on STX3B as a molecular on-off switch might be due to the opening of STX3B through phosphorylation of its N-peptide.

Whether or not STX3B is palmitoylated in the retinal ribbon synapses is unknown but the presence of cysteines in its TMD hints at a potential for palmitoylation. On the other hand, it is also possible that the E259 might render the TMD of STX3B inadequate for palmitoylation as in the case of STX1A^K260E^. This raises the possibility of inhibition of spontaneous vesicle fusion in retinal ribbon synapses by the non-palmitoylated TMD of STX3B. This is interesting as the retinal ribbon synapses operate in an essentially different manner compared to the conventional synapses as they predominantly rely on the tonic fusion but not on AP-driven phasic fusion of the SVs. Thus, JMD dependent inhibition of the TMD palmitoylation of STX3B and therefore the blockage of the spontaneous release even at the expense of the full capacity of Ca^2+^-evoked release might be beneficial for a ribbon synapse to enhance the signal-to-noise ratio of the tonic neurotransmitter release.

Our chimeric analysis of the TMD of different syntaxins also show that besides the regulation of the efficacy of SV fusion by the TMD of STX1A, spontaneous SV fusion is further modulated by the C-terminal half of STX1A’s SNARE domain suggesting a direct involvement of STX1A in SV clamp. Which intermolecular interactions regulate the SNARE domain mediated clamping on spontaneous release is not clear, as known interactions of STX1A with the modulatory proteins SYT1 and CPX are shown to be carried out through the N-terminal half of its SNARE domain (Chen et al., 2002; Zhou et al., 2015; Zhou et al., 2017). Nevertheless, this suggests STX1A contributes to the regulation of spontaneous release through two distinct mechanisms, one is inhibitory through its C-terminal half of SNARE domain and the other is facilitatory through the palmitoylation of its TMD.

## Material and Methods

### Animal maintenance and generation of mouse lines

All procedures for animal maintenance and experiments were in accordance with the regulations of and approved by the animal welfare committee of Charité-Universitätsmedizin and the Berlin state government Agency for Health and Social Services under license number T0220/09. The generation of STX1-null mouse line was described previously (Arancillo et al., 2013; Vardar et al., 2016).

### Neuronal cultures and lentiviral constructs

Hippocampal neurons were obtained from mice of either sex at postnatal day (P) 0–2 and seeded on the priorly prepared continental or micro-island astrocyte cultures as described previously (Vardar et al., 2016; Xue et al., 2007). The neuronal cultures were then incubated for 13-20 DIV in Neurobasal^™^-A supplemented with B-27 (Invitrogen), 50 IU/ml penicillin and 50 μg/ml streptomycin at 37°C before experimental procedures. Neuronal cultures were transduced with lentiviral particles at DIV 1-3. Lentiviral particles were provided by the Viral Core Facility (VCF) of the Charité-Universitätsmedizin, Berlin, and were prepared as previously described (Vardar et al., 2016). The cDNAs of mouse STX1A (NM_016801), STX3A (NM_152220), and STX3B (NM_001025307) were cloned in frame after an NLS-GFP-P2A sequence within the FUGW shuttle vector (Lois et al., 2002) in which the ubiquitin promoter was replaced by the human synapsin 1 promoter (f(syn)w). The improved *Cre* recombinase (iCre) cDNA was C-terminally fused to NLS-RFP-P2A. SynGCamp6f was generated analogous to synGCamp2 (Herman et al., 2014), by fusing GCamp6f (Chen et al., 2013) to the C terminus of synaptophysin and within the f(syn)w shuttle vector (Grauel et al., 2016).

### Western Blot

The lysates were obtained from DIV13-16 high-density neuronal cultures cultivated in 35 mm culture dishes. The neurons were lysed in 200 μl lysis buffer containing 50 mm Tris/HCl, pH 7.9, 150 mm NaCl, 5 mm EDTA, 1% Triton X-100, 0.5% sodium deoxycholate, 1% Nonidet P-40, and 1 tablet of Complete Protease Inhibitor (Roche) for 30 min on ice. Equal amounts of solubilized proteins were loaded in 12% SDS-PAGE and subsequently transferred to nitrocellulose membranes. The membranes were subjected to the following primary antibodies overnight at 4°C according to the experiment: mouse monoclonal anti-β-tubulin III (1:5000; Sigma) or mouse monoclonal anti-actin (1:4000) as internal controls, mouse monoclonal anti-STX1A (1:10.000; Synaptic Systems), mouse monoclonal anti-SNAP25 (1:10.000; Synaptic Systems), and rabbit polyclonal anti-STX3 (1:1000; Synaptic Systems). Horseradish peroxidase (HRP)-conjugated goat secondary antibodies (Jackson ImmunoResearch) were applied for 1 h at room temperature and detected with ECL Plus Western Blotting Detection Reagents (GE Healthcare Biosciences) in Fusion FX7 image and analytics system (Vilber Lourmat).

### Immunocytochemistry

The high-density cultured hippocampal neurons cultivated on 12 mm culture dishes were fixed with 4% paraformaldehyde (PFA) in 0.1 M phosphate-buffered saline, PH 7.4, for 10 min at DIV13-16. The neurons were then permeabilized with 0.1% Tween^®^-20 in PBS (PBST) for 45 min at room temperature (RT) and then blocked with 5% normal goat serum (NGS) in PBST. Primary antibodies, guinea pig polyclonal anti-Bassoon (1:1000; Synaptic Systems) and rabbit polyclonal anti-STX3 (1:1000; Synaptic Systems),were applied overnight at 4°C. Subsequently, secondary antibodies, rhodamine red donkey anti-guinea pig IgG (1:500; Jackson ImmunoResearch) and A647 donkey anti-rabbit IgG (1:500; Jackson ImmunoResearch) were applied for 1 h at RT in the dark. The coverslips were mounted on glass slides with Mowiol^®^ mounting agent (Sigma-Aldrich). The images were acquired with an Olympus IX81 epifluorescence-microscope with MicroMax 1300YHS camera using MetaMorph software (Molecular Devices). Exposure times of excitations were kept constant for each wavelength throughout the images obtained from individual cultures. Data were analyzed offline with ImageJ as previously described (Vardar et al., 2016). Sample size estimation was done as previously published (Vardar et al., 2016).

### Acyl-Biotin Exchange (ABE) Method

2-3 x 10^6^ neurons were cultivated on 100 mm culture dishes coated with astrocyte culture and transduced at DIV 1 with *Cre* recombinase in combination with STX1A^WT^, STX1A^K260E^ or STX1A^CVCV^. All STX1A constructs used for ABE method were N-terminally tagged with FLAG epitope.

For the biotinylation of the proteins, the ABE method was applied as previously described (Brigidi and Bamji, 2013). Briefly, the lysates were obtained at DIV 13-16 in lysis buffer, pH 7.2, including 50 mM NEM that was freshly dissolved in EtOH at the day of the experiment. After incubation on ice for 30 min, the lysates were centrifuged and the supernatant was transferred into a fresh tube. 50 μl of anti-FLAG magnetic beads (Sigma) were then added into the lysates for overnight incubation on a rotator at 4°C. Next day, the supernatant was discarded and the anti-FLAG magnetic beads were nutated in lysis buffer, pH 7.2, including 10 mM NEM for 10 min on ice. The beads were then washed thrice with lysis buffer, pH 7.2,. The beads were then resuspended with 1 ml lysis buffer, pH 7.2 and separated into half. The supernatant was discarded and one sample from each group was treated with 500 μl lysis buffer, pH 7.2 including 1 M HAM and the other sample without HAM. The HAM solution was freshly prepared at the day of the experiment. The HAM cleavage proceeded for 1 h on a rotator at RT.

After HAM cleavage, the STX1A bound anti-FLAG magnetic beads were washed thrice in lysis buffer, pH 7.2, and once in lysis buffer, pH 6.2. The beads were then resuspended in lysis buffer, pH 6.2, including 1 μM Biotin-BMCC for 1 h on a rotator at 4°C. Biotin-BMCC covalently binds to the thiol groups on cysteine residues exposed after the HAM driven cleavage of the palmitates. The supernatant was discarded and the beads were washed thrice in lysis buffer, pH 7.2. The beads were then resuspended in 30 μl 1X PBS and 1X SDS-PAGE loading buffer. After incubation at 95°C for 5 min, the supernatant was loaded on an SDS-PAGE and subsequently transferred to a nitrocellulose membrane. The membranes were treated with HRP-conjugated streptavidin antibody (ThermoFisher) for 1 h at RT. The chemiluminescence was detected in Fusion FX7 image and analytics system (Vilber Lourmat) after treatment of the membrane with ECL Plus Western Blotting Detection Reagents (GE Healthcare Biosciences).

### Electrophysiology

Whole cell patch-clamp recordings were performed on glutamatergic autaptic hippocampal neurons at DIV 14–20 at RT with a Multiclamp 700B amplifier and an Axon Digidata 1550B digitizer controlled by Clampex 10.0 software (both from Molecular Devices). The recordings were analyzed offline using Axograph X Version 1.7.5 (Axograph Scientific).

Prior to recordings, the transduction of the neurons was verified by RFP and GFP fluorescence. Membrane capacitance and series resistance were compensated by 70% and only the recordings with a series resistance smaller than 10 MΩ were used for further recordings. Data were sampled at 10 kHz and filtered by low-pass Bessel filter at 3 kHz. The standard extracellular solution was applied with a fast perfusion system (1–2 ml/min) and contained the following: 140 mM NaCl, 2.4 mM KCl, 10 mM HEPES, 10 mM glucose, 2 mM CaCl_2_, and 4 mM MgCl_2_ (300 mOsm; pH 7.4). Borosilicate glass patch pipettes were pulled with a multistep puller, yielding a final tip resistance of 2–5 MΩ when filled with KCl based intracellular solution containing the following: 136 mM KCl, 17.8 mM HEPES, 1 mM EGTA, 4.6 mM MgCl_2_, 4 mM ATP-Na_2_, 0.3 mM GTP-Na_2_, 12 mM creatine phosphate and 50 U/ml phosphocreatine kinase (300 mOsm; pH 7.4).

The neurons were clamped at −70 mV in steady state. To evoke EPSCs, the neurons were depolarized to 0 mV for 2 ms. The size of the RRP was determined by a 5 s application of 500 mM sucrose in standard external solution (Rosenmund and Stevens, 1996) and the total charge transfer was calculated as the integral of the transient current. Fusogenicity measurement was conducted by application of 250 mM sucrose solution for 10 s and calculation of the ratio of the charge transfer of the transient current over RRP. Spontaneous release was determined by monitoring mEPSCs for 30 - 60 s at −70 mV. To correct false positive events, mEPSCs were recorded in the presence of 3 μM AMPA receptor antagonist NBQX (Tocris Bioscience) in standard external solution.

Sample size estimation was done as previously published (Rosenmund and Stevens, 1996).

### SynGcamp6f-imaging

Imaging experiments were performed at DIV 13-16 on autapses in response to a single stimulus and trains of stimuli at 10 Hz as described previously for SynGcamp2-imaging (Herman et al., 2014). Images were acquired using a 490-nm LED system (pE2; CoolLED) at a 5 Hz sampling rate with 25 ms of exposure time. The acquired images were analyzed offline using ImageJ (National Institute of Health), Axograph X (Axograph), and Prism 8 (Graph-Pad; San Diego, CA). Sample size estimation was done as previously published (Herman et al., 2014).

### Statistical analysis

Data in violin graphs present single observations (points), median and the quartiles (lines). Data in x-y plots present means ± SEM. All data were tested for normality with Kolmogorov-Smirnov test. Data from two groups with normal or non-parametric distribution were subjected to Student’s two-tailed t-test or Mann-Whitney non-parametric test, respectively. Data from more than two groups were subjected to Kruskal-Wallis followed by Dunn’s *post hoc* test when at least one group showed a non-parametric distribution. For data in which all the groups showed a parametric distribution, one-way ANOVA test followed by Tukey’s *post hoc* test was applied. All the tests were run with GraphPad Prism 8.3 and all the statistical data are summarized in corresponding source data tables.

## Acknowledgements

We thank the Charité Viral Core facility, Katja Pötschke, and Bettina Brokowski for virus production, Berit Söhl-Kielczynski and Heike Lerch for technical asssistance and to all the Rosenmund Lab members for the discussions. This work was supported by the German Research Council (DFG) grants 388271549, 399894546, 436260754, 278001972.

---

**Figure 2 – Supplement 1:**
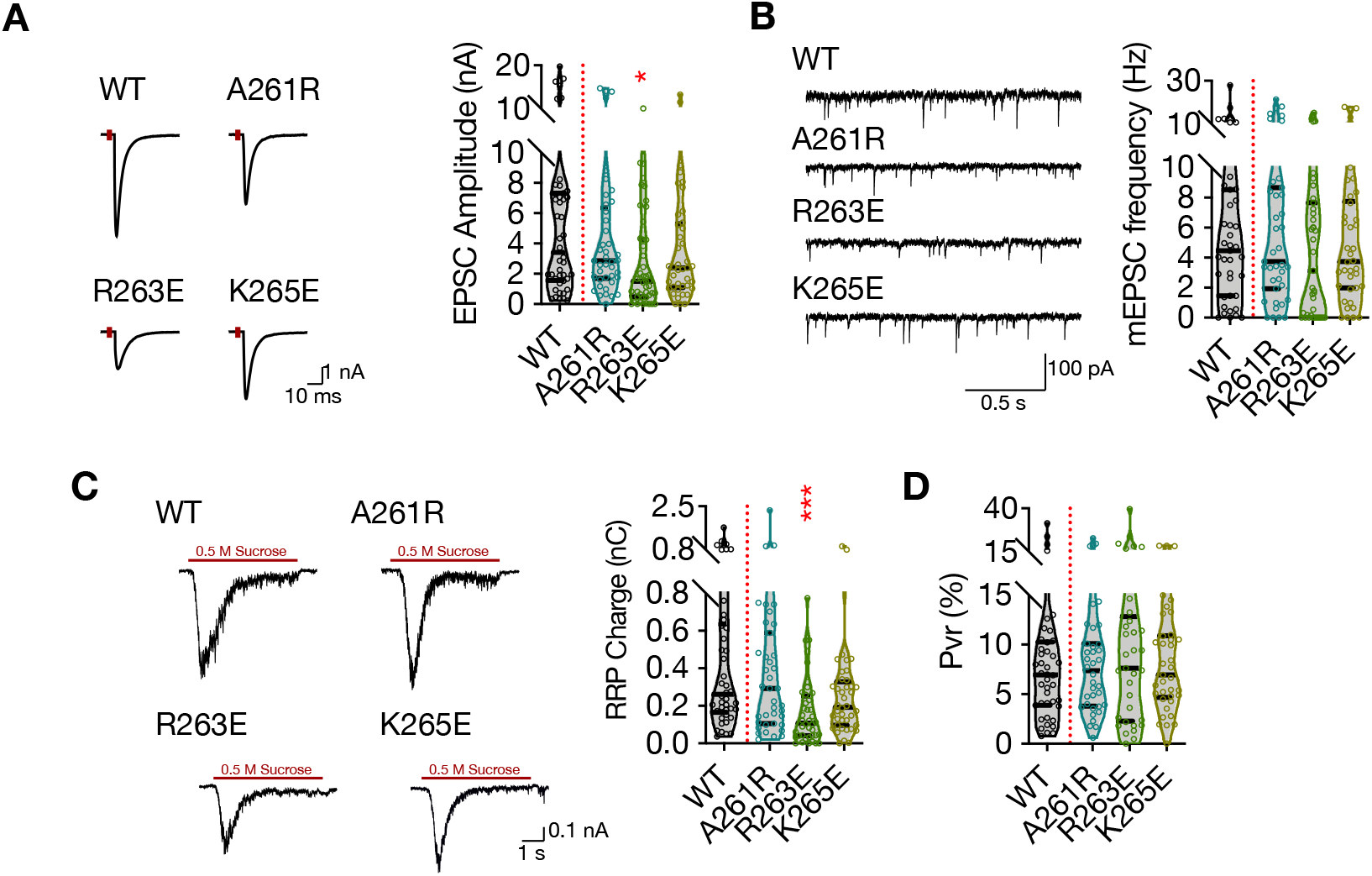
Charge reversal mutations in STX1A’s JMD manifest position specific effects on different modes of neurotransmitter release. A. Example traces (left) and quantification of the amplitude (right) of EPSCs obtained from hippocampal autaptic STX1-null neurons rescued either with STX1A^WT^, STX1A^A261R^, STX1A^R263E^, or STX1A^K264E^. B. Example traces (left) and quantification of the frequency (right) of mEPSCs recorded from the same neurons as in (A). C. Example traces (left) and quantification of RRP recorded from the same neurons as in (A). D. Quantification of Pvr recorded from the same neurons as in (A). Data information: The artifacts are blanked in example traces in (A) and (C). The example traces in (B) were filtered at 1 kHz. In (A–D), data points represent single observations, the violin bars represent the distribution of the data with lines showing the median and the quartiles. Red annotations (stars) on the graphs show the significance comparisons to STX1A^WT^. Nonparametric Kruskal-Wallis test followed by Dunn’s *post hoc* test was applied to data in (A-D); *p ≤ 0.05, ***p ≤ 0.001. The numerical values are summarized in source data.

**Figure 4 – Supplement 1:**
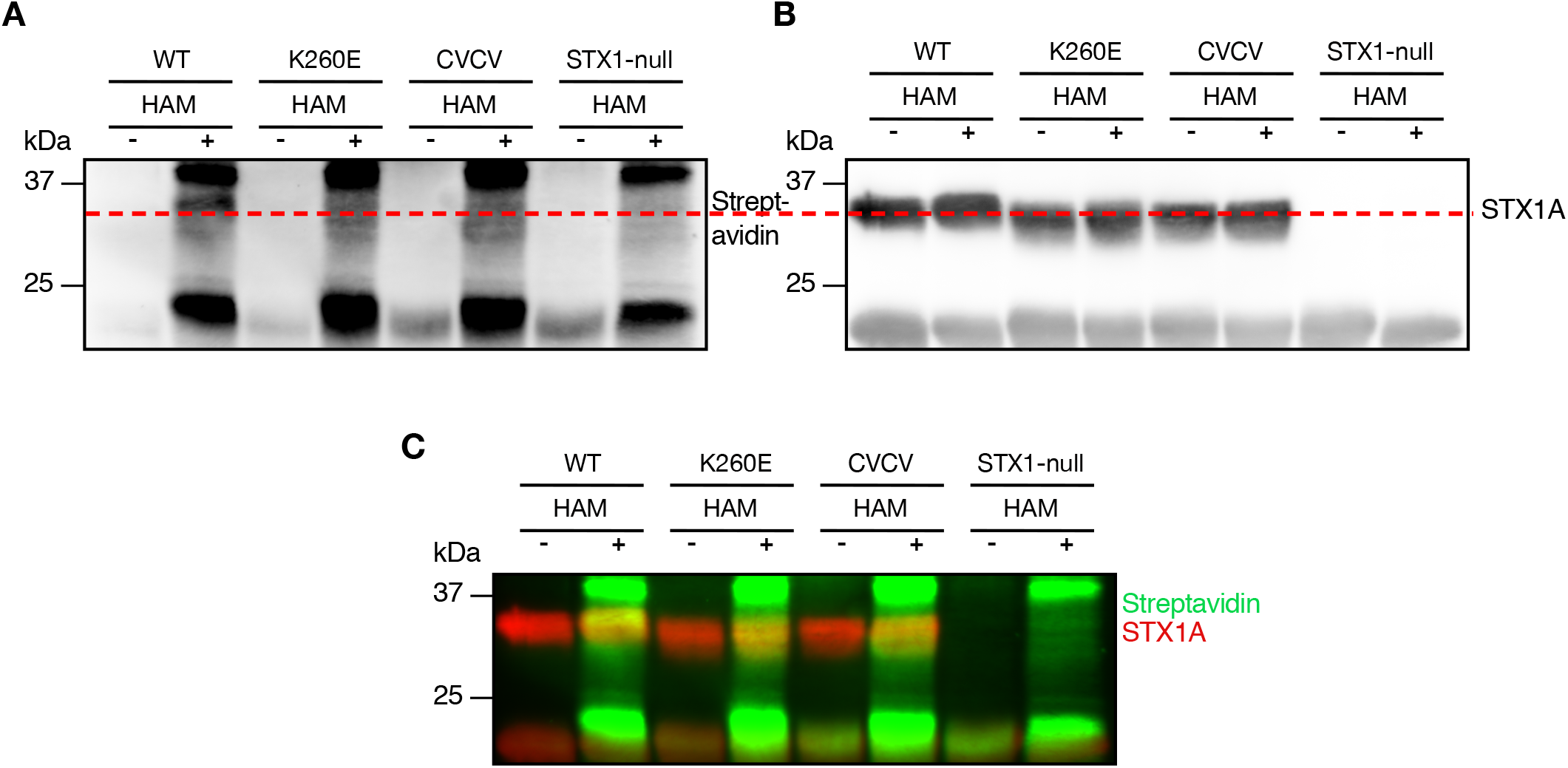
K260E mutation in STX1A’s JMD leads to loss of palmitoylation of its TMD. A. Palmitoylation was assessed by ABE method. The samples were probed with HRP-streptavidin antibody. B. The HRP-streptavidin antibody was stripped and the samples were further probed with anti-STX1A antibody. The red dashed line indicates that the biotin that covalently bound to the free thiols after HAM cleavage in (A) is detected at the position of STX1A in (B). C. Superimposition of the biotin and STX1A detection.

**Figure 5 – Supplement 1:**
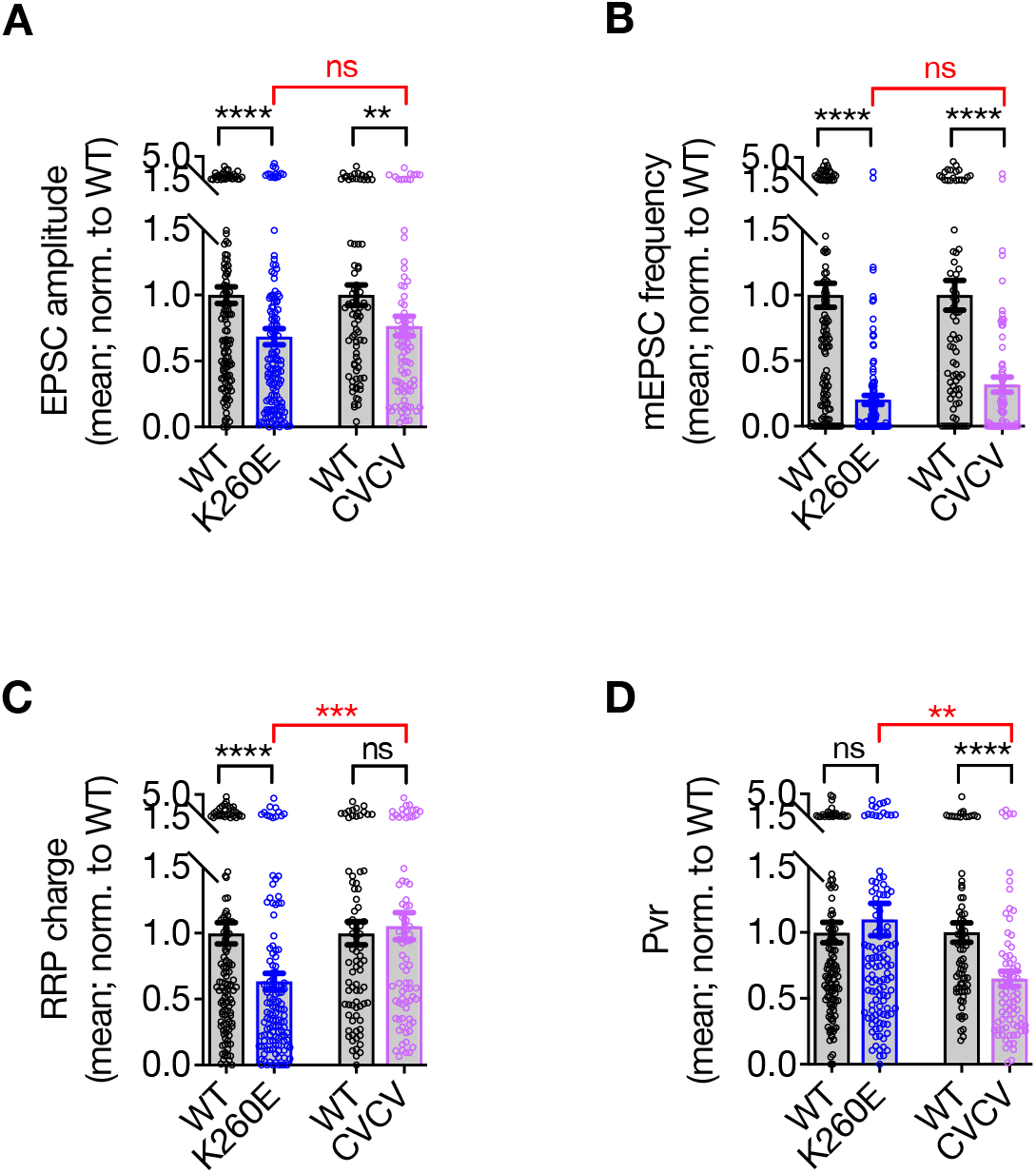
Comparison of the STX1A^K260E^ and STX1A^CVCV^ phenotypes by the pooled data of the normalized values for each culture. A. The EPSC amplitude values were normalized to STX1A^WT^ in each individual culture and pooled together. B. The mEPSC frequency values were normalized to STX1A^WT^ in each individual culture and pooled together. C. The RRP values were normalized to STX1A^WT^ in each individual culture and pooled together. D. The Pvr values were normalized to STX1A^WT^ in each individual culture and pooled together. Data information: Data points represent single observations and bar graphs represent mean ± SEM. Black annotations (stars and ns) on the graphs show the significance comparisons to STX1A^WT^ based on nonparametric Mann-Whitney test Red annotations (stars and ns) on the graphs show the significance comparisons between STX1A^K260E^ and STX1A^CVCV^ based on nonparametric Kruskal-Wallis test followed by Dunn’s *post hoc* test; **p ≤ 0.01, ***p ≤ 0.001, ****p ≤ 0.0001. The numerical values are summarized in source data.

**Figure 8 – Supplement 1:**
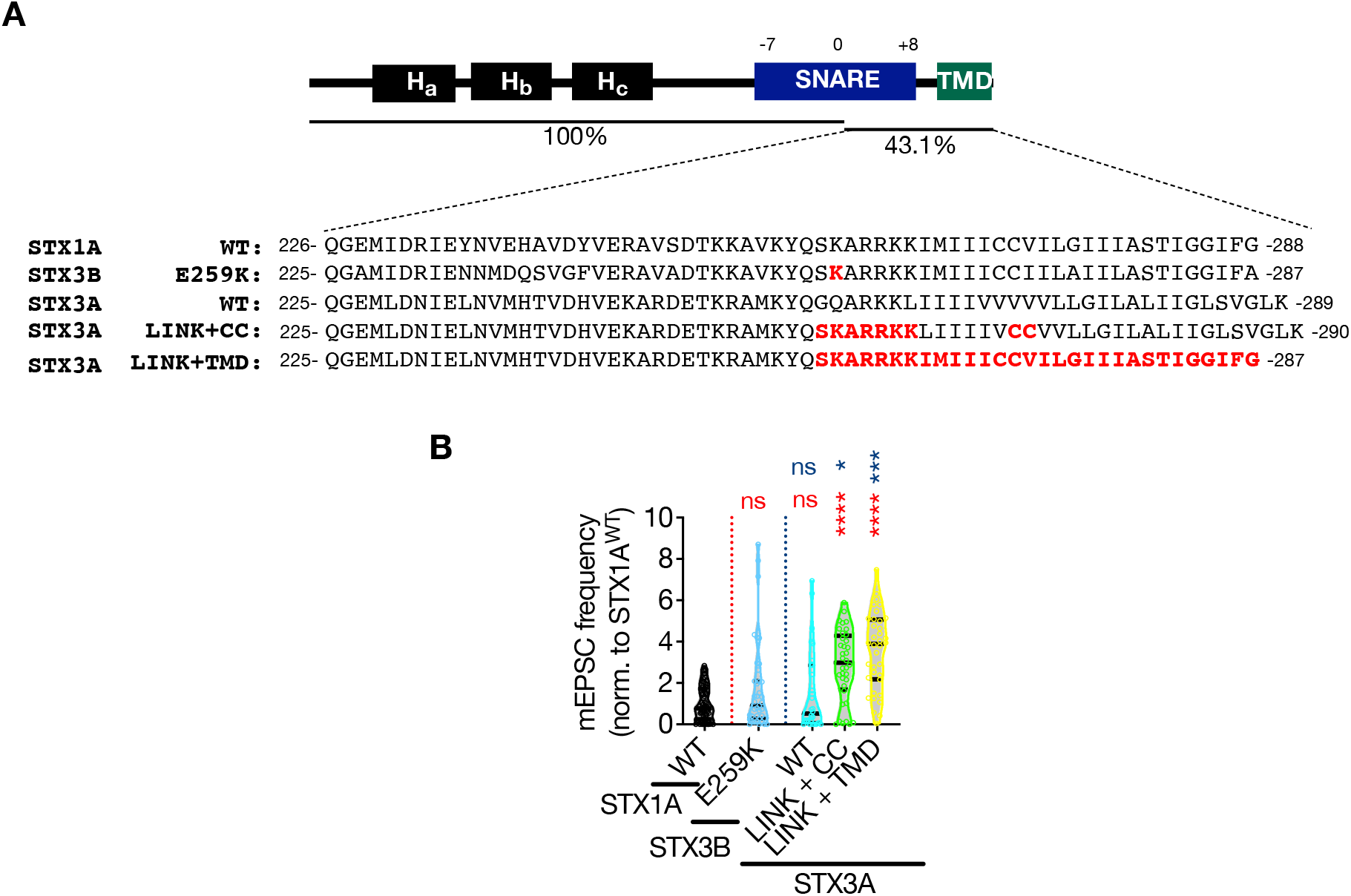
C-terminal half of STX1A’s SNARE domain clamps spontaneous neurotransmitter release. A. Sequence alignment of the JMD and TMD regions of STX1A^WT^, STX3B^E259K^, STX3A^WT^, STX3A^LINK+CC^, and STX3A^LINK+TMR^. B. The mEPSC frequency values were normalized to STX1A^WT^ in each individual culture and pooled together. Data information: Data points represent single observations and the violin bars represent the distribution of the data with lines showing the median and the quartiles. Red and blue annotations (stars and ns) on the graphs show the significance comparisons to STX1A^WT^ and STX3B^E259K^, respectively. Nonparametric Kruskal-Wallis test followed by Dunn’s *post hoc* test was applied to data in (C-F); *p ≤ 0.05, ***p ≤ 0.001, ****p ≤ 0.0001. The numerical values are summarized in source data.

## Source Data file titles and legends

**Figure 1 – Source Data 1**

**Table 1:** Quantification of the neurotransmitter release parameters of STX1-null neurons lentivirally transduced with STX1A^WT^ or with STX1A JMD elongation mutants

**Figure 2 – Source Data 1**

**Table 2:** Quantification of the neurotransmitter release parameters of STX1-null neurons lentivirally transduced with STX1A^WT^ or with STX1A JMD charge reversal mutants

**Figure 2 – Supplement 1 - Source Data 1**

**Table 3:** Quantification of the neurotransmitter release parameters of STX1-null neurons lentivirally transduced with STX1A^WT^ or with STX1A JMD charge reversal mutants −2

**Figure 3 – Source Data 1**

**Table 4:** Quantification of the neurotransmitter release parameters of STX1-null neurons lentivirally transduced with STX1A^WT^ or with STX1A JMD charge reversal mutants −3

**Figure 3 – Source Data 2**

Whole SDS-PAGE images represented in Figure 3A and Figure 3D

**Figure 4 – Source Data 1**

**Table 5:** Quantification of the neurotransmitter release parameters of STX1-null neurons lentivirally transduced with STX1A^WT^ or with STX1A palmitoylation mutans

**Figure 4 – Source Data 2**

Whole SDS-PAGE image represented in Figure 4B

**Figure 4 – Supplement 1 - Source Data 1**

Whole SDS-PAGE images represented in Figure 4C and Figure 4 – Supplement 1

**Figure 5 – Source Data 1**

**Table 6:** Quantification of the neurotransmitter release parameters of STX1-null neurons transduced either with STX1A^WT^, STX1A^K260E^, STX1A^CVCV^, or STX1A^K260E+CVCV^

**Figure 5 – Supplement 1 - Source Data 1**

**Table 7:** Quantification of the neurotransmitter release parameters of STX1A^K260E^ and STX1A^CVCV^ neurons as normalized to the values recorded from STX1A^WT^ neurons

**Figure 6 – Source Data 1**

**Table 8:** Quantification of the neurotransmitter release parameters of STX1-null neurons lentivirally transduced with STX1A^WT^ or K260 mutants

**Figure 6 – Source Data 2**

Whole SDS-PAGE image represented in Figure 6B

**Figure 7 – Source Data 1**

**Table 9:** Quantification of the neurotransmitter release parameters of STX1-null neurons lentivirally transduced with STX1A^WT^ or with STX3B^WT^ or E259K mutant

**Figure 8 – Source Data 1**

**Table 10:** Quantification of the neurotransmitter release parameters of STX1-null neurons lentivirally transduced with STX1A^WT^ or with STX3A^WT^ or mutants

**Figure 8 – Source Data 2**

Whole SDS-PAGE images represented in Figure 7B and Figure 8B

**Figure 8 – Supplement 1 - Source Data 1**

**Table 11:** Quantification of the neurotransmitter release parameters of STX3A and STX3B neurons as normalized to the values recorded from STX1AWT neurons

**Figure 9 – Source Data 1**

**Table 12:** Quantification of the presynaptic Ca^2+^-influx in STX1-null neurons transduced either with STX1A^WT^, STX3B^WT^ or mutant, or STX3A^WT^ or mutants

